# Mapping sites of carboxymethyllysine modification on proteins reveals its consequences for proteostasis and cell proliferation

**DOI:** 10.1101/2020.10.16.342311

**Authors:** Simone Di Sanzo, Katrin Spengler, Anja Leheis, Joanna M. Kirkpatrick, Theresa L. Rändler, Tim Baldensperger, Luca Parca, Christian Marx, Zhao-Qi Wang, Marcus A. Glomb, Alessandro Ori, Regine Heller

**Author notes:** These authors contributed equally to this work (Simone Di Sanzo, Katrin Spengler). These authors jointly supervised this work (Alessandro Ori, Regine Heller).

## Abstract

Posttranslational mechanisms play a key role in modifying the abundance and function of cellular proteins. Among these, modification by advanced glycation end products (AGEs) has been shown to accumulate during aging and age-associated diseases but specific protein targets and functional consequences remain largely unexplored. Here, we devised a proteomic strategy to identify specific sites of carboxymethyllysine (CML) modification, one of the most abundant AGEs. We identified over 1000 sites of CML modification in mouse and primary human cells treated with the glycating agent glyoxal. By using quantitative proteomics, we found that protein glycation triggers a proteotoxic response and directly affects the protein degradation machinery. We show that glyoxal induces cell cycle perturbation in primary endothelial cells and that CML modification reduces acetylation of tubulins and impairs microtubule dynamics. Our data demonstrate the relevance of AGE modification for cellular function and pinpoint specific protein networks that might become compromised during aging.

**Highlights:**

- A peptide enrichment strategy allows mapping of CML modification in cells and tissues
- CML modification competes with ubiquitination or acetylation of lysines
- Glyoxal treatment destabilizes the 26S proteasome
- Glyoxal arrests cell cycle and impairs microtubule dynamics via altering the tubulin code

## Introduction

Advanced glycation end products (AGEs) are generated via a non-enzymatic glycation of proteins initiated by the reaction of glucose, fructose or highly reactive dicarbonyls such as methylglyoxal or glyoxal with amino acids (Brownlee, 1995). Glycation is enhanced when glucose levels are high, when dicarbonyls deriving from glycolytic intermediates or lipid peroxidation accumulate (Nigro et al., 2019), or when dicarbonyl detoxification by glyoxalases such as glyoxalase 1 (GLO1) and protein deglycase DJ-1 is low (Nigro et al., 2017). AGE formation may alter structure and function of the targeted proteins and, in addition, AGEs may elicit their effects as ligands of the pro-inflammatory receptor for advanced glycation end products (RAGE) (Teissier and Boulanger, 2019). One of the most abundant AGEs *in vivo* is N(6)-carboxymethyllysine (CML), whose formation is triggered, among other routes, by glyoxal (Brings et al., 2017). CML is known to be chemically stable, to accumulate in human tissues in diabetes, atherosclerosis, neurodegeneration and aging, and thus to be a biomarker of aging (Delgado-Andrade, 2016).

A causal relationship between the buildup of AGEs and individual disease is still under debate, but studies in which AGE levels were modulated via the GLO1 system, provided first evidence. Overexpression of GLO1 increases lifespan in *C. elegans* (Morcos et al., 2008) and reduces endothelial dysfunction in diabetic mice (Brouwers et al., 2014), while knockdown of GLO1 mimics diabetic nephropathy in non-diabetic mice (Giacco et al., 2014). Glycation of extracellular proteins such as hemoglobin HbA1c (Shapiro et al., 1980) or collagen (Avery and Bailey, 2005) is well described, but reports on glycation of intracellular proteins are still scarce. Examples of those are histones (Ansari et al., 2018; Baldensperger et al., 2020; Galligan et al., 2018; Guedes et al., 2011; Mir et al., 2014; Zheng et al., 2019; Zheng et al., 2020), mitochondrial proteins (Hamelin et al., 2007; Rosca et al., 2005; Wang et al., 2009), the 20S proteasome (Queisser et al., 2010), enzymes involved in energy production (Snow et al., 2007), small heat shock proteins (Oya-Ito et al., 2006; Schalkwijk et al., 2006; Sudnitsyna and Gusev, 2017) and the sodium channel Nav1.8 (Bierhaus et al., 2012). Glycation is increasingly seen as a driver of metabolic disease and aging, and it may elicit specific effects by targeting signaling proteins (Chaudhuri et al., 2018; Kold-Christensen and Johannsen, 2020). In this context, glycation of the ryanodine receptor associated with successive Ca^2+^ leakage and mitochondrial damage (Ruiz-Meana et al., 2019), reversible inhibitory glycation of nuclear factor erythroid 2-related factor 2 (Nrf2) (Sanghvi et al., 2019), and methylglyoxal-induced dimerization of Kelch-like ECH-associated protein 1 (KEAP1) with subsequent activation of the KEAP1/Nrf2 transcriptional program (Bollong et al., 2018) have been reported. The latter may play a role in the upregulation of defense systems, such as GLO1 and the ubiquitin-proteasome system (UPS), and may be involved in the hormetic effect of methylglyoxal (Ravichandran et al., 2018). In contrast to earlier assumptions of irreversible AGE formation, recent studies suggested the possibility of deglycation via DJ-1 or protein arginine deiminase 4 (Zheng et al., 2019; Zheng et al., 2020).

A global characterization of the targeted proteins and pathways, which may significantly contribute to manifestation of metabolic diseases and aging, is a prerequisite to further understand pathophysiological effects of AGEs, like CML. Here, we developed a proteomic workflow based on selective enrichment of CML-modified peptides coupled to mass spectrometry for identification and quantification. We applied this approach to two cellular models (mouse embryonic fibroblasts (MEF) and human umbilical vein endothelial cells (HUVEC)), and to tissues of young and chronologically aged mice. This strategy allowed us to identify specific sites of CML modification. We show that CML modification of proteins competes with ubiquitination or acetylation, and that it destabilizes components of the UPS. CML accumulation in primary endothelial cells led to inhibition of proliferation, which was due to both altered expression of cell cycle regulators, and glycation of tubulins associated with impaired microtubule dynamics.

## Results

### An enrichment strategy for the identification of CML-modified peptides

Aiming to identify sites of CML modification on proteins, we developed a proteomic workflow employing an antibody-based enrichment of CML-modified peptides coupled to mass spectrometry (CMLpepIP). We applied CMLpepIP to MEF and HUVEC treated with different concentrations of glyoxal, a cell-permeable dialdehyde (Fig. 1A). In both cell types, glyoxal treatment led to a dose-dependent increase of total CML levels, as verified by immunoblot (Fig. 1B), and amino acid absolute quantification using liquid chromatography mass spectrometry (LC-MS) (Fig. 1C). To assess the efficiency of our enrichment protocol, we compared the fraction of peptide spectrum matches (PSM) assigned to CML-containing peptides between elutions from the enriched and control samples. When CMLpepIP was applied to glyoxal-treated cells, we achieved up to 7 times more PSM from CML-containing peptides as compared to controls. With a lower concentration of glyoxal, the gain of modified peptides was reduced and, in any case, the fraction of PSM assigned to CML peptides never exceeded 5% (Fig. 1D). This indicates that CMLpepIP enables reproducible (Fig. S1A and S1B), but only partial, enrichment of CML-modified peptides, especially in high complexity samples where CML peptides occur at low concentration (<1%).

**Figure 1.**
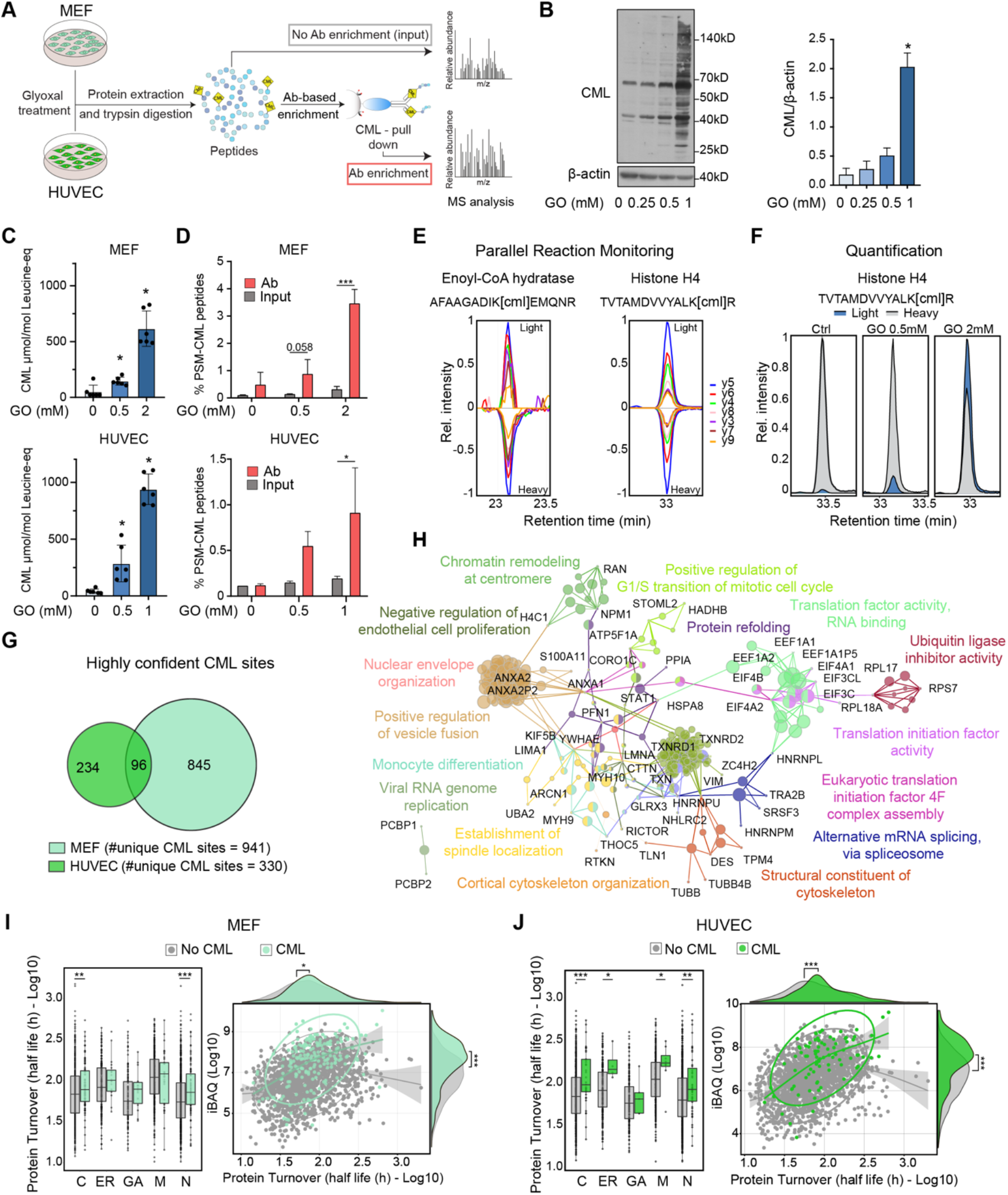
Antibody-based enrichment of CML-modified peptides. **A.** Workflow for the identification of CML-modified peptides. Ab: antibody **B.** Immunoblot (left) and densitometry-based quantification (right) for CML-modified proteins from HUVEC treated with glyoxal (GO) for 48 h. n=3, * p<0.05 vs. control, one-way repeated measurement ANOVA corrected using Holm-Šidák method. **C.** Quantification of total CML levels in MEF and HUVEC treated with GO for 8 h or 48 h, respectively. n=6, * p<0.05 vs. control, one-way ANOVA using Geisser-Greenhouse correction. **D.** Percentage of peptide spectrum matches (PSM) containing CML modification among different conditions. n=3 (MEF), n=2 (HUVEC), * p<0.05, *** p<0.001. Multiple t-test corrected using Holm-Šidák method. **E.** Validation of CML-modified peptides by parallel reaction monitoring (PRM) using heavy spike-in peptides. **F.** Quantification of a CML-modified peptide from histone H4 by PRM in MEF treated with GO for 24h. **G.** Venn diagram showing the overlap between unique highly confident CML sites identified in MEF and HUVEC. **H.** Enrichment of Gene Ontology biological processes among CML sites identified in both HUVEC and MEF. The enrichment was performed using the Cytoscape App ClueGO. **I-J.** Left, protein turnover of CML-modified proteins grouped according to their subcellular localization, as defined by Gene Ontology annotation. Turnover data were taken from (Mathieson et al., 2018). C: cytoplasm; ER: endoplasmic reticulum; GA: Golgi apparatus; M: mitochondria; N: nucleus. Right, scatterplot comparing protein abundance (iBAQ scores, averages of n=3 (MEF), n=2 (HUVEC)) and protein turnover of CML-modified proteins. * p<0.05, ** p<0.01, *** p<0.001, Wilcoxon Rank Sum test with continuity correction. Related to Fig. S1 and Table S1.

Using CMLpepIP, we identified 995 unique CML sites in MEF and 451 in HUVEC of which 941, and 330 were deemed ‘highly confident’, respectively (see Methods for applied cut-offs) (Fig. S1A and Table S1). Overlapping sets of CML sites were reproducibly identified across conditions and independent experiments (Fig. S1B and S1C). To validate the identified CML peptides, we performed parallel reaction monitoring (PRM) measurements using isotopically labeled spike-in peptide standards derived from our list of identified CML sites (Table S1). Co-elution of endogenous (light) and standard (heavy) peptides validated the identified CML-modified peptides, and it enabled quantitation relative to the spike-in standard across conditions that recapitulated the absolute levels of CML measured by LC-MS (Fig. 1E,F).

A subset of conserved CML sites was found to be modified in both MEF and HUVEC (Fig. 1G). These sites localized on proteins involved in translation, RNA binding and chromatin remodeling at the centromere as well as on structural constituents of the cytoskeleton and mitochondrial proteins (Fig. 1H and Table S1). Since long-lived proteins are postulated to be the main targets of AGEs (Gkogkolou and Bohm, 2012; Smuda et al., 2015; Verzijl et al., 2000), we used published turnover data (Mathieson et al., 2018) and investigated the turnover distribution for modified and non-modified proteins in MEF and HUVEC. In both cell types, proteins with slower turnover were more likely to be affected by CML modification, independently of their subcellular localization (Fig. 1I left - J left), indicating slow turnover as an important determinant of CML modification. Although we were able to detect CML sites across the entire dynamic range of identified proteins in both cell types (Fig. S1D), we noted that protein abundance correlated with CML modifications, with high abundant proteins being more often detected as modified (Fig. 1I right - J right).

Taken together, CMLpepIP enabled us to investigate the CML-modified proteome and to identify specific and conserved targets of AGE-modification in two different primary culture systems. Lower turnover and high abundance appear to be key factors in driving CML modification *in vitro*.

### Sites of CML modification in mouse organs

Next, we applied CMLpepIP to investigate protein targets of CML *in vivo*. We analyzed heart, kidney and liver from young (3-4 months) and geriatric C57BL/6J mice (26-33 months), and, in parallel, age-related changes of protein abundance using quantitative MS (Fig. 2A). The selected organs showed a higher basal level of total CML content compared to, e.g., brain (Fig. S2A). We identified 257, 186 and 52 unique CML sites in heart, kidney and liver, respectively, of which 190, 142 and 34 were deemed ‘highly confident’, respectively (Table S2). Using PRM with spike-in heavy peptides, we validated 16 (89%) out of those 18 CML sites, for which we had successfully developed PRM assays (Fig. S2B-C). CML appeared to target proteins in an organ-specific manner (Fig. 2B). The affected proteins participate in biological processes related to energy production in the heart, cytoskeleton in the kidney, and detoxification in the liver (Fig. 2C). Similar to what we observed in cultured cells (Fig. 1I,J), CML sites in the heart appeared to occur preferentially in proteins with slow turnover (Fig. S2D). Since CML modification occurs on lysines and these residues are targets for other posttranslational modifications (PTM), we assessed the overlap of CML sites and other known PTM. We found that approximately one third of the identified CML sites (97 out of 334 unique highly confident sites) occur on residues that are known to be modified by other PTM, primarily ubiquitination and acetylation (Fig. 2D). These data suggest that modification by CML might interfere with biological processes by impeding other PTM to occur on specific residues.

**Figure 2.**
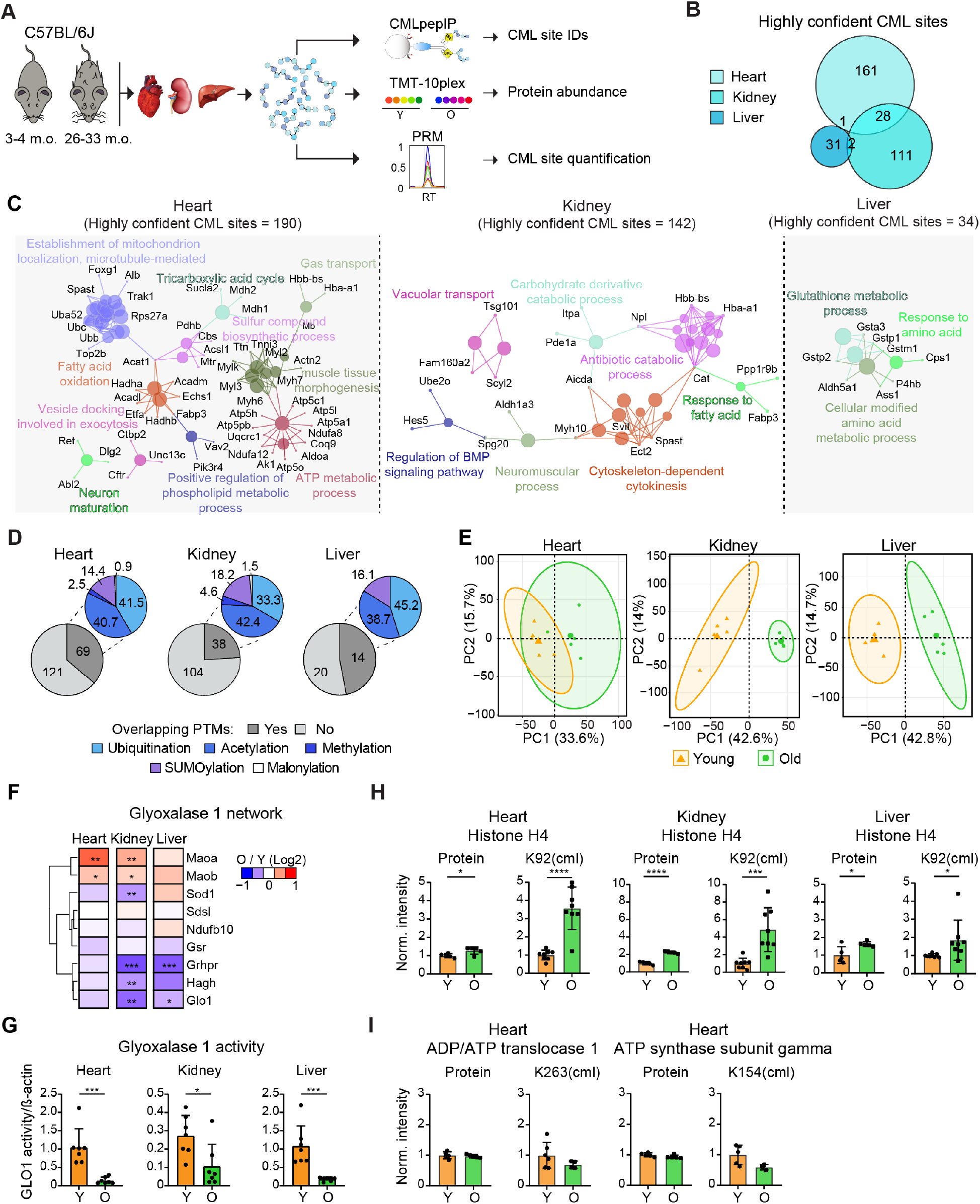
The CML-modified proteome in mouse organs. **A.** Workflow for the analysis performed on mouse tissues from young (Y) and old (O) mice. Tandem Mass Tag (TMT) was used for monitoring protein abundance, CMLpepIP for CML site identification and PRM for CML site quantification. **B.** Venn diagram of highly confident CML sites identified in heart, kidney and liver. **C.** Enrichment of Gene Ontology biological processes among CML sites identified from heart, kidney and liver. The enrichment was performed using the Cytoscape App ClueGO. **D.** Overlap between CML sites and other PTM. The gray scale pie charts indicate the number of CML sites overlapping with known PTM. The colored pie charts indicate which PTM class overlap with CML sites (shown as percentages). Known PTM were obtained from Minguez et al., 2015; Hornbeck et al., 2015). **E.** Principal component analysis of proteome data from young and old mice (n=5 per age group). The smaller dots represent individual samples and the larger dots the centroids of each age-matched group. Ellipses represent 95% confidence intervals. The percentage of variance explained by the first two principle components (PC) axes is reported in the axis titles. **F.** Effect of aging on the abundance of proteins part of the glyoxalase 1 (GLO1) network. The network was extracted from https://string-db.org using input GLO1. n=5, * adj. p<0.05, ** adj. p<0.01, *** adj. p<0.001, t-test with Benjamini-Hochberg correction for multiple testing, as implemented in *limma*. **G.** GLO1 activity tested on heart, kidney and liver lysates from young (Y) and old (O) mice. n=7, * p<0.05, *** p<0.001, unpaired t-test, parametric, two-tailed. **H-I.** Normalized protein intensity of histone H4, ADP/ATP translocase type 1 and ATP synthase subunit γ from TMT experiment (Protein - left panel) and intensity of CML-modified peptides from PRM (right panel). n=5 for protein abundance, n=8 for CML-modified peptides, * p<0.05, *** p<0.001, **** p<0.0001, unpaired t-test. Related to Fig. S2 and Table S2.

We next investigated age-related changes of protein abundance in the same tissues using Tandem Mass Tag (TMT)-10plex labeling (Fig. 2A). Principal component analysis could clearly distinguish two different clusters formed by young and old mice for all the three organs (Fig. 2E). By applying gene set enrichment analysis, we identified pathways affected by protein abundance changes in aged organs (Fig. S2E). Among these, a pathway related to AGE signaling was enriched with aging both in heart and kidney. This pathway included components of the fibrotic process such as several types of collagen I and IV, but also proteins mediating signaling downstream of extracellular AGEs via RAGE such as NFkB1, MAPK9 and AKT1. In addition, we specifically queried the data for proteins involved in dicarbonyl detoxification via GLO1 and found a general trend for a decreased abundance of the components of this network in all organs isolated from geriatric mice (Fig. 2F). We validated the functional relevance of decreased GLO1 protein levels by confirming a decrease of its activity in old organs (Fig. 2G).

Inspired by this and by the well-established increase of total AGE levels in old tissue (Chaudhuri et al., 2018; Delgado-Andrade, 2016; Nigro et al., 2019), we tested whether we could detect an age-dependent elevation of CML modification at specific sites. Using PRM assays on total tissue lysates, we demonstrated an age-dependent raise in the level of CML modification of histone H4 at position K92 that exceeded the abundance change observed at the protein level in heart and kidney (Fig. 2H). However, no age-related changes in the levels of two other CML sites, i.e., K263 of adenine nucleotide translocase type 1 and K154 of ATP synthase subunit γ, were observed (Fig. 2I).

Together, these data show that the targets of AGEs are organ-specific, and that the levels of CML modification increase with aging in a site-specific manner, likely as a consequence of reduced levels and activity of dicarbonyl-detoxifying enzymes in old organs.

### CML modification directly targets the ubiquitin-proteasome system

To better understand the consequences of increased CML modifications in tissues, we performed experiments to characterize cellular functions in response to glyoxal treatment. Since we observed that some of the CML targets are proteins involved in protein quality control (Fig. S3A), we hypothesized that accumulation of glyoxal impairs proteostasis and triggers cellular responses to counteract proteotoxic stress. To address this question, we applied a data-independent acquisition (DIA) method to monitor proteome-wide changes of protein abundance in MEF treated with different glyoxal concentration for 8 h and 24 h (Fig. 3A and S3B). Short treatment (8 h) with 0.5 mM and 2 mM glyoxal induced only few significant changes of protein abundance, however, a longer treatment (24 h) triggered a major proteome response that was dose-dependent (Fig. 3B). At this time point, the abundance of CML-modified proteins was significantly increased (Fig. 3C), while KEGG pathways related to the UPS, especially ubiquitin-conjugating enzymes (E2), were decreased (Fig. 3D and S3C).

**Figure 3.**
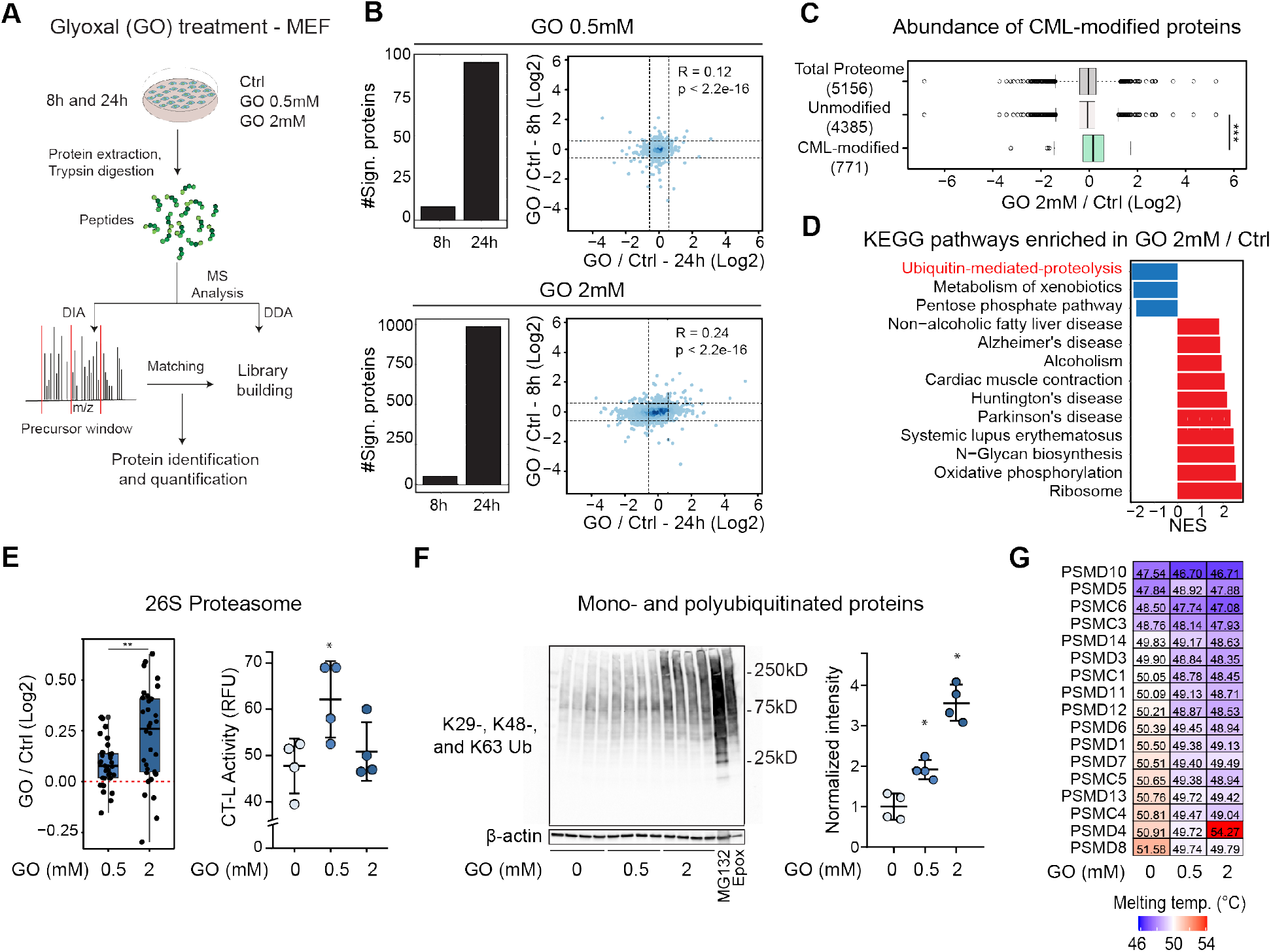
The proteome response to glyoxal (GO) treatment. **A.** Workflow for the analysis of proteome changes induced in MEF by GO. **B.** Left, number of significant proteins (absolute log2 fold change>0.58 and q<0.05) after 8 and 24 h GO treatment. Right, scatterplots comparing the log2 fold changes induced by GO at 8 or 24 h. Dashed lines represent the log2 fold change cut-offs used (±0.58). n=5 (8 h), n=4 (24 h). **C.** Distribution of protein fold changes for 2 mM GO vs. control (Ctrl) after 24 h of treatment. Average values of n=4 were compared, *** p<0.001, Wilcoxon rank sum test continuity correction with alternative two-sided. **D.** Gene Set Enrichment Analysis (GSEA) for KEGG pathways based on protein fold changes induced by 2 mM GO after 24 h of treatment. Normalized enrichment score (NES) indicates pathways enriched among proteins that increase (red) or decrease (blue) upon GO treatment (FDR<0.05). All the proteins quantified were ranked according to their log2 fold change and used as input for GSEA. **E.** Boxplot of fold changes for members of the 26S proteasome (left) and proteasome chymotrypsin-like (CT-L) activity (right) in MEF treated with GO for 24 h compared to Ctrl. n=4, ** p<0.01, Wilcoxon Rank Sum test with continuity correction. **F.** Immunoblot for mono- and polyubiquitinated proteins from MEF lysates treated with GO for 24 h. Lysates from cells treated with proteasome inhibitors MG132 (20 μM for 6h) or epoxomicin (Epox, 10 nM for 96h) were loaded as positive controls. n=4, * p<0.05, one-way ANOVA using Geisser-Greenhouse correction. **G.** Heatmap representing the melting point temperature of members of 26S proteasome identified in all conditions obtained using thermal proteome profiling. Related to Fig. S3 and Table S3.

Interestingly, we found a significant increase of components of the 26S proteasome induced by glyoxal treatment in a dose-dependent manner in both MEF and HUVEC (Fig. 3E left and S3D). However, while 0.5 mM glyoxal also increased proteasomal activity in MEF, the induction of 26S proteasome by 2 mM glyoxal was not followed by a corresponding increase of activity (Fig. 3E right). In parallel, we observed a dose-dependent accumulation of ubiquitinated proteins 24 h after glyoxal treatment (Fig. 3F). We speculated that the lack of proteasome activity increase at high doses of glyoxal might be due to a functional impairment of the proteasome itself. To test this, we evaluated the thermal stability of proteasomal proteins in MEF treated with glyoxal using thermal proteome profiling (Franken et al., 2015). This analysis revealed that glyoxal treatment negatively impacted the thermal stability of most of the 26S proteasome subunits in a dose-dependent fashion (Fig. 3G), while the overall thermal stability of proteins remained comparable across the tested conditions (Fig. S3E and Table S3). In addition, we identified a subset of proteasomal proteins and other members of the UPS, including ubiquitin itself, to be directly CML-modified both in cells treated with glyoxal and in organs (Fig. S3A).

Taken together these data describe a specific and dose-dependent proteome response to glyoxal exposure. This is characterized by accumulation of CML-modified proteins and an induction of compensatory mechanisms including the UPS. The UPS may become saturated at higher glyoxal concentration due to the accumulation of CML-modified and ubiquitinated proteins, and to a destabilization of the 26S proteasome.

### Glyoxal inhibits proliferation of primary human endothelial cells by inducing cell cycle arrest and inhibiting microtubule dynamics

In endothelial cells, glyoxal-induced CML modifications occurred mainly in proteins of the nucleus, the cytoskeleton and the mitochondria (Fig. 4A and Table S4), while glyoxal-initiated changes in protein abundance comprised an upregulation of stress response proteins, e.g., heme oxygenase 1, and a downregulation of processes related to proliferation and growth (Fig. 4B). In particular, several factors involved in DNA replication were downregulated. This was also reflected on a functional level since glyoxal led to an inhibition of endothelial cell proliferation as demonstrated by reduced cell numbers and lower incorporation of BrdU in the presence of serum or after stimulation with basic fibroblast growth factor (bFGF) (Fig. 4C,D). Furthermore, angiogenic sprouting in response to vascular endothelial growth factor (VEGF) was impaired (Fig. 4E). Glyoxal-treated cells did not show signs of apoptosis (Fig. S4A).

**Figure 4.**
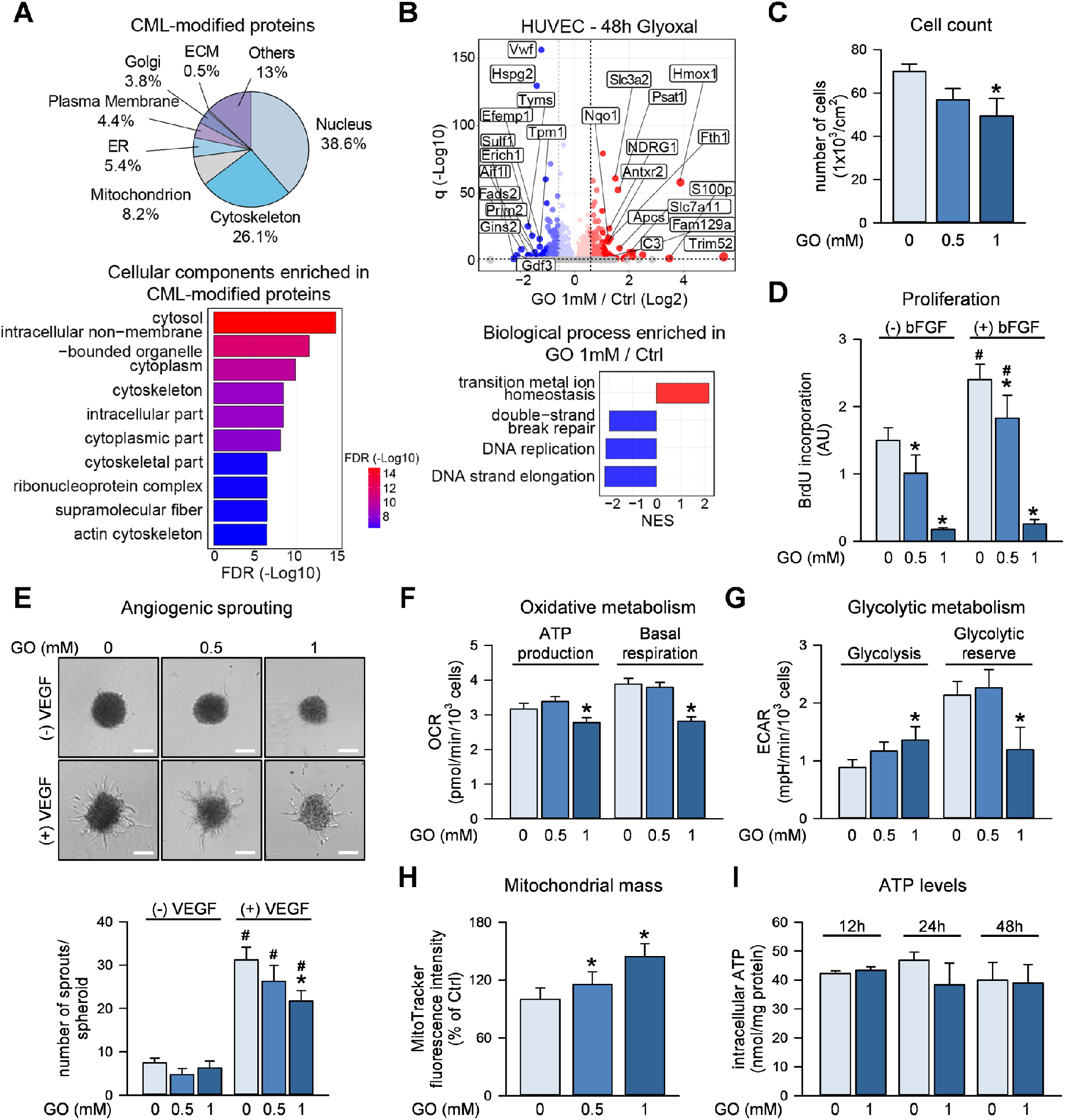
Glyoxal (GO) impairs the proliferation of HUVEC. **A.** Upper panel: CML-modified proteins annotated to different cellular compartments according to Gene Ontology annotation after GO treatment (1 mM, 48 h). Lower panel: Gene Ontology cellular component terms enriched in CML-modified proteins. False Discovery Rate (FDR) <0.05. **B.** Upper panel: volcano plot depicting proteins that significantly increase (red) or decrease (blue) abundance upon GO treatment (1 mM, 48 h) or remain unchanged (gray). Horizontal dashed line indicates a significance cut-off of q<0.05 and vertical dashed lines an absolute fold change (log2)>0.58. Lower panel: gene set enrichment analysis for Gene Ontology biological process terms based on protein fold changes. Terms enriched among increased (red) or decreased (blue) proteins are shown. FDR<0.05; NES: normalized enrichment score. **C-H**. HUVEC were treated with GO for 48 h. **C**. Cell numbers were determined. n=5. **D**. Cells were stimulated with bFGF (50 ng/ml, 24 h) and BrdU incorporation was measured. n=5. **E**. Spheroids were generated, embedded and stimulated with VEGF (50 ng/ml, 24 h). Representative pictures and sprout numbers per spheroid are shown. Scale bar=100 μm. n=4. **F-G**. Oxygen consumption rate (OCR) (**F**) and extracellular acidification rates (ECAR) (**G**) were measured via Seahorse technology and mitochondrial and glycolytic parameters calculated. n=6. **H**. Cells were stained with MitoTracker and analyzed by flow cytometry. n=5. **I**. HUVEC were treated with 1 mM GO and intracellular ATP levels were measured in cell extracts. n=3. **C-I**. Data are represented as mean ± SEM. Statistical significance was analyzed using one-way or two-way repeated measurement ANOVA corrected using Holm-Šidák method. * p<0.05 vs. control, # p<0.05 vs. respective non-bFGF or non-VEGF-treated sample. Related to Fig. S4 and Table S4.

Since mitochondrial proteins, e.g., ATP synthase subunit γ, were found to be CML-modified (Fig. 1H), we asked whether alterations in cellular energy production would underlie the observed inhibition of cell proliferation. Seahorse analyses revealed a decrease in basal respiration and mitochondrial ATP production in HUVEC treated with glyoxal (1 mM, 48 h) (Fig. 4F and Fig. S4B) as well as an upregulation of glycolysis (Fig. 4G and Fig. S4C). The latter, together with an increased mitochondrial biogenesis (Fig. 4H), likely compensated the reduction of mitochondrial ATP production, since total cellular ATP was not altered (Fig. 4I). These data show that the cellular energy homeostasis was maintained despite the exposure of cells to dicarbonyl stress and that reduced cell proliferation was not due to metabolic limitations.

To further clarify the mechanisms underlying inhibition of cell proliferation by glyoxal, we analyzed its effect on the cell cycle. Applying a triple staining method, we found that treating endothelial cells with glyoxal for 24 h triggered an arrest in G0/G1 and G2 phases, while the percentage of cells in the S and M phases was significantly decreased (Fig. 5A). Some of these alterations, especially the decrease in mitosis, were already seen after 4 h of glyoxal treatment, although at a lower degree. In accordance with the lower entry of cells into S and M phases, we observed a decrease of the cell cycle markers cyclin A (S phase) and phosphorylated H3(S10) (p-H3(S10), M phase) (Fig. S5A). In addition, tracking cell proliferation dynamics revealed a slower proliferation in response to the high dose of glyoxal (Fig. 5B).

**Figure 5.**
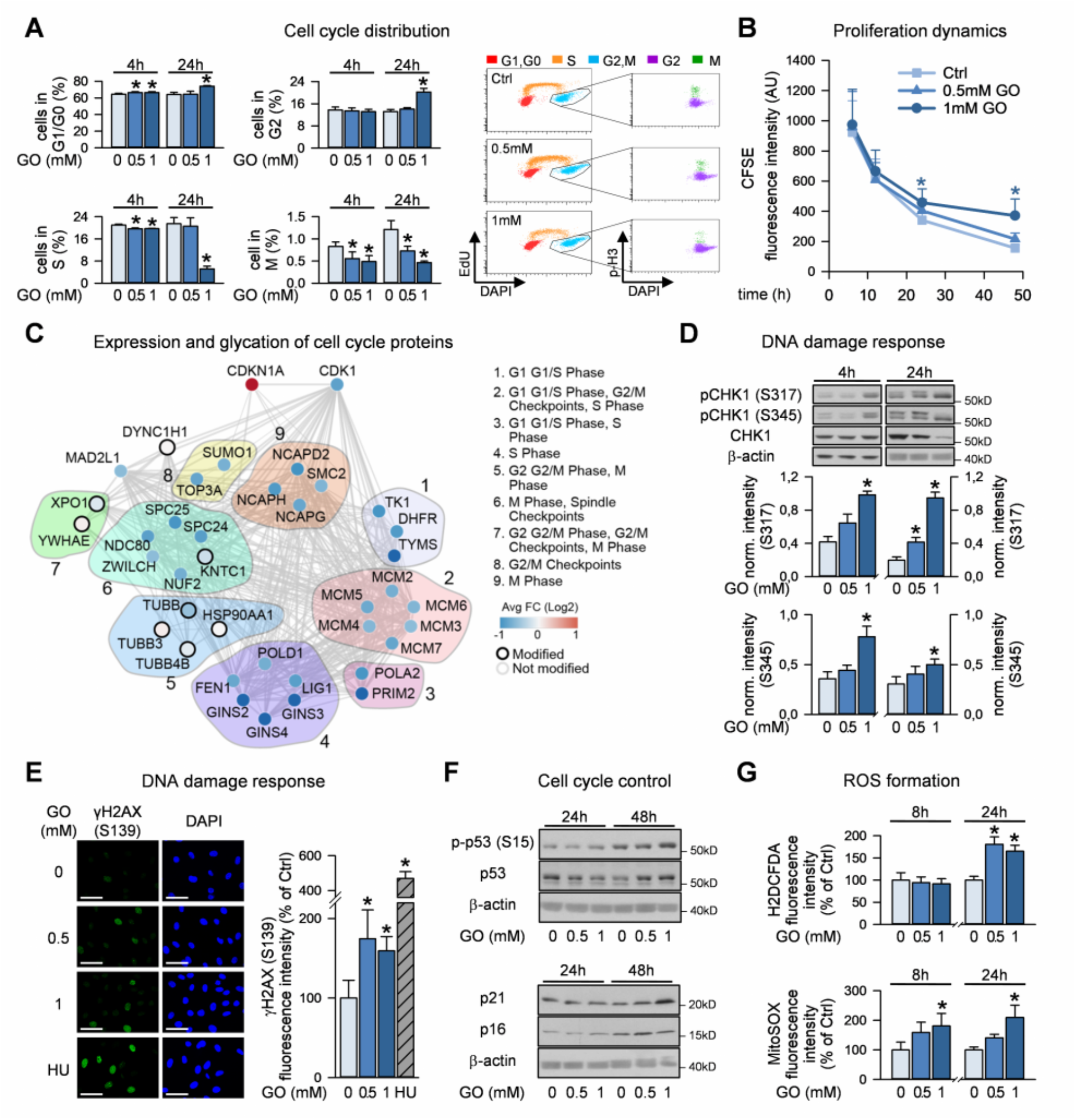
Perturbation of the cell cycle induced by glyoxal (GO). **A**. HUVEC were treated with GO as indicated and subjected to cell cycle analysis applying a triple staining method (EdU, p-H3 (S10), DAPI). Ratio of cells in the respective cell cycle phases (left) and representative dot plots (right) are shown. n=4. **B**. HUVEC were stained with carboxyfluorescein succinimidyl ester (CFSE) and treated with GO. CFSE staining was quantified by flow cytometry. n=4. **C**. Cell cycle proteins were chosen according to Reactome annotation divided by their specific role in each phase. Only proteins with significantly altered expression (absolute fold change>0.58 and q<0.05) or modified by CML are shown. Each dot represents the protein fold change and the black edge signifies proven CML modification. **D**. GO-treated HUVEC were lysed and subjected to immunoblot analysis. n=5 (4 h), n=6 (24 h). **E**. Following GO treatment of HUVEC (8h), immunofluorescence staining of γH2AX (S139) and DAPI was performed. Hydroxyurea (HU, 2 mM, 1 h) served as positive control. Representative pictures and quantification of γH2AX (S139)-positive nuclei are shown. Scale bar=50 μm. n=3. **F**. HUVEC were treated with GO, lysed and subjected to immunoblot analysis. Densitometric quantification is shown in figure S5B. n=6. **G**. HUVEC were treated with GO and intracellular (H2DCFDA) and mitochondrial ROS (MitoSOX) were measured by flow cytometry. n=4. Statistical significance was analyzed using one-way or two-way repeated measurement ANOVA corrected using Holm-Šidák method. * p<0.05 vs. respective control. Related to Fig. S5 and Table S5.

We hypothesized that the observed cell cycle arrest induced by glyoxal was due to altered expression of cell cycle regulators plus to specific glycation of proteins involved in cell cycle control, which were both identified in glyoxal-treated cells (Fig. 5C). Interestingly, alteration of protein expression was mainly related to proteins involved in the regulation of S and G1 phases, while glycation was mostly observed in proteins controlling G2 and M phases, for instance, in tubulin a and β chains (Fig. 5C). Strikingly, 8 out of 10 subunits of the replicative helicase MCM2-7/GINS/CDC45 and several factors involved in lagging strand DNA synthesis such as subunits of DNA polymerases α and δ, FEN1 and DNA ligase 1 were downregulated, whereas the cyclin-dependent kinase inhibitor 1A (CDKN1A)/p21 was upregulated.

One pathway through which the expression of cell cycle regulators is affected is the DNA damage response. Glyoxal triggered phosphorylation of the checkpoint kinase 1 (CHK1) and led to an increased percentage of cells with γH2AX foci (Fig. 5D,E). As a consequence, and in line with the proteomics data, the p53/p21 pathway was upregulated by glyoxal (Fig. 5F and S5B). Immunoblotting experiments showed that especially with the higher dose of glyoxal the abundance and, additionally, the phosphorylation of p53 as well as the expression of its target CDKN1A/p21 increased. The expression of another inhibitor of cyclin-dependent kinases, p16, was enhanced, too (Fig. 5F). We also found that after treatment with 1 mM glyoxal a small proportion of cells (~12%) acquired a senescent phenotype, as shown by positive staining for senescence-associated beta-galactosidase (SA-β-Gal) (Fig. S5C). This may be related to persistent dicarbonyl and oxidative stress since glyoxal led to increased mitochondrial and cytosolic formation of reactive oxygen species (ROS) (Fig. 5G).

To understand how, on the other hand, glycation of proteins may affect cell cycle progression, we focused on tubulins, which exhibited CML modification of different lysines upon glyoxal treatment and in tissues from old mice (Fig. 6A and S6). CML-modified tubulin showed more strongly structured tubulin filaments when compared to tubulin in control cells (Fig. 6B), and an increased stability against the depolymerizing agent nocodazole (Fig. 6C). These data point to an impairment of microtubule dynamics upon glyoxal treatment, which may affect mitosis and contribute to the observed reduction of cell proliferation. To check this, we compared mitosis in glyoxal-treated and control cells independent from G1 and G1/S transition. We performed a double thymidine block to synchronize cells in early S phase, treated them with glyoxal thereafter and monitored time-dependent expression of the mitotic marker p-H3(S10) in immunoblotting experiments. In control cells, p-H3(S10) signals were detected between 16 h and 18 h after releasing the cell cycle block, while glyoxal-treated cells showed reduced intensity of p-H3(S10) signals indicating a lower number of mitotic cells (Fig. 6D).

**Figure 6.**
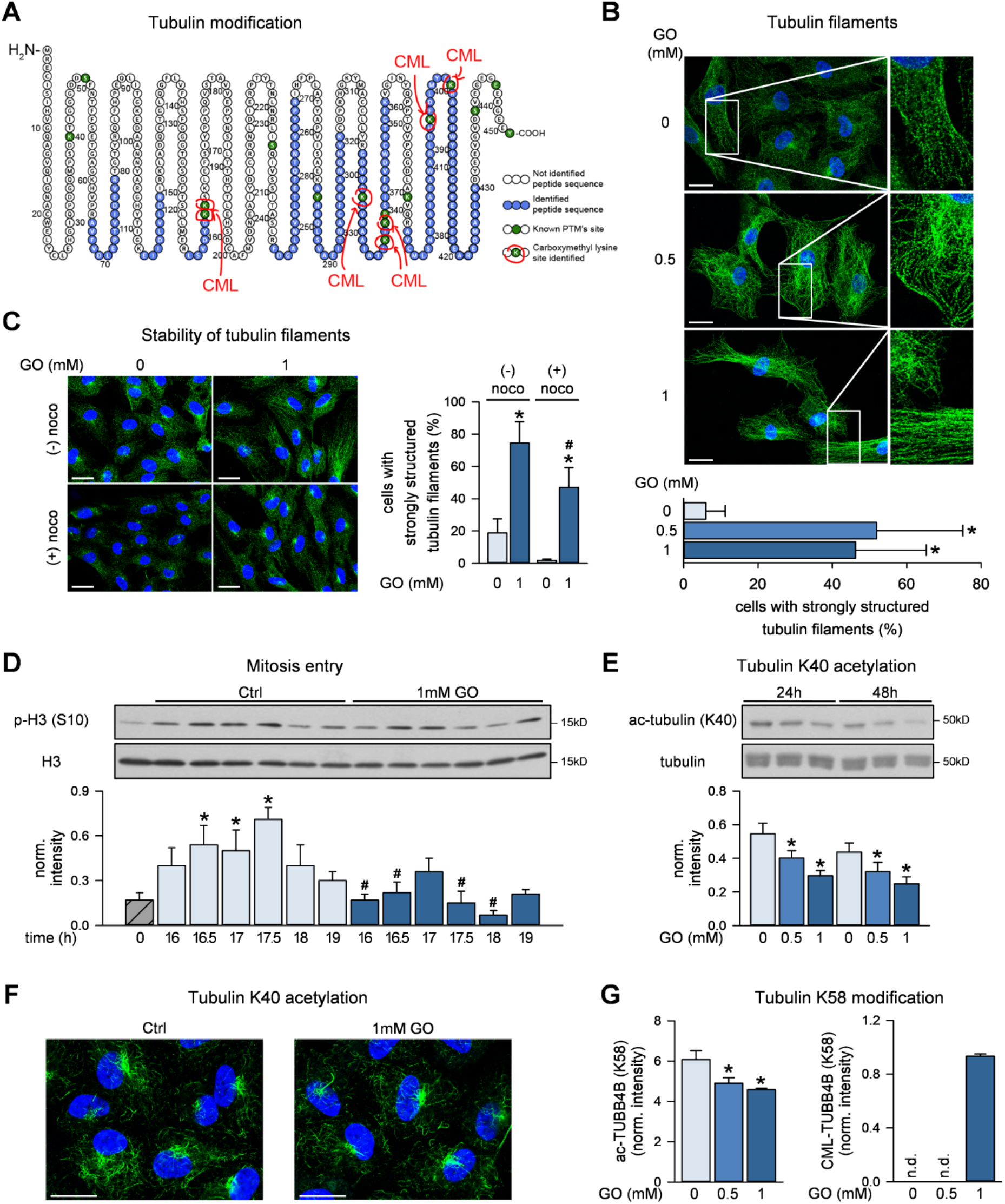
Glyoxal (GO) affects dynamics and post-translational modification of tubulins. **A**. Scheme of detected CML modification sites for tubulin α-1B chain. **B**. HUVEC were treated with GO for 48 h and stained against α/β-tubulin and DAPI. Representative immunofluorescence pictures (upper panel) and the percentage of cells with strongly structured tubulin filaments per high-power field (lower panel) are shown. Scale bar=20 μm. n=4. **C**. HUVEC were treated with GO (1 mM, 48 h). 10 μM nocodazole was added for 2 min and α/β-tubulin and DAPI were stained. Representative pictures (left) and the percentage of cells with strongly structured tubulin filaments per high-power field (right) are shown. Scale bar=20 μm. n=5. **D**. HUVEC were synchronized via a double thymidine block and treated with GO subsequent to releasing the block. Cells were lysed and subjected to immunoblot analysis n=5. **E**. HUVEC were treated with GO, lysed and subjected to immunoblot analysis. n=6. **F**. HUVEC were treated with GO for 24 h and stained against ac-tubulin (K40) and DAPI. A representative picture out of n=3 is shown. **G**. HUVEC were treated with GO for 48 h. Tubulin modifications were detected by mass spectrometry after respective enrichment of CML-modified or acetylated lysines. Barplot shows the intensity of tubulin β acetylated (left panel) or CML-modified at K58 (right panel). The peptide used for the barplot is _INVYYNEATGGK*YVPR_. Statistical significance was analyzed using one-way or two-way repeated measurement ANOVA corrected using Holm-Šidák method. * p<0.05 vs. control, # p<0.05 vs. respective non-nocodazole-treated cells (C) or vs. respective non-GO-treated cells (D). Related to Fig. S6.

CML modification of tubulin may directly affect microtubule dynamics but may also compete with other lysine modifications, thereby affecting tubulin functionality. Since the lysine residues of tubulin, which are modified by CML, are also known to be acetylated (Sadoul and Khochbin, 2016) (Fig. 6A), we hypothesized that CML modification may impede tubulin acetylation. Indeed, glyoxal treatment of HUVEC led to reduced tubulin acetylation at lysine 40 (K40) (Fig. 6E-F), which is known to protect microtubules from mechanical damage and to facilitate self-repair (Portran et al., 2017; Xu et al., 2017). Of note, we were able to demonstrate a direct competition of CML modification and acetylation at K58 of the tubulin β-4B chain in glyoxal-treated cells. Glyoxal induced a significant CML modification of K58 and at the same time, K58 acetylation was reduced (Fig. 6G). Together, these data suggest that CML modification of tubulin is likely to have an impact on acetylation and may thereby affect its interactions with microtubule-binding proteins and regulatory enzymes.

## Discussion

The impairment of protein homeostasis is a hallmark of aging and age-associated diseases (Hipp et al., 2019; Ori et al., 2015; Santos and Lindner, 2017). While age-related changes in protein abundance and turnover have been described across tissues and in different species, less is known about the extent and potential functional impact of PTM. Here, we focused on CML modification of proteins, a non-enzymatic PTM belonging to the class of AGEs (Delgado-Andrade, 2016). We developed an antibody-based method for the enrichment of CML-modified peptides coupled to mass spectrometry for identification and quantification. The mapping of CML sites together with studying the total proteome in cultured cells and aging mouse organs allowed us to link CML modifications to cellular functions and to better understand how these processes may contribute to aging and age-associated metabolic diseases.

We validated our method in cultured cells treated with the glycating agent glyoxal and identified over 1000 specific sites of CML modification. CML occurs more often in high abundant proteins that display slower turnover across different cellular compartments. This corresponds to previous studies showing that global AGEs accumulate particularly in tissues characterized by slow protein turnover, such as crystallin lens, cartilage and skin (Gkogkolou and Bohm, 2012; Smuda et al., 2015; Verzijl et al., 2000). Interestingly though, we detected CML-modified peptides across proteins that spanned four orders of magnitude of protein abundance and, conversely, we did not identify CML modifications for all of the most abundant proteins. Keeping in mind potential biases due to limited sensitivity and specific protein sequences not amenable to enzymatic digestion that are inherent to any proteomic analysis, our data suggest the existence of a subset of proteins that are more prone to CML modification than others.

We extended our analysis to primary organs collected from young and old mice and showed that our enrichment strategy is able to identify hundreds of physiologically occurring CML sites, some of these previously reported in an independent study in the mouse heart (Ruiz-Meana et al., 2019). The modified proteins were enriched for mitochondrial proteins, histones, cytoskeletal proteins and enzymes involved in detoxification. We additionally developed targeted proteomics assays that enable the quantification of a subset of CML sites in tissues without prior peptide enrichment. These analyses revealed that the level of CML modification for some sites, i.e., histone H4-K92 (corresponding to K91 in the mature protein), increased with aging in different organs, while others remained constant. Thus, while global AGE levels increase with age, preferential modification of certain proteins occurs, as previously suggested in plants (Bilova et al., 2017), illustrating the need for targeted investigation of AGE modification sites. Interestingly, increased CML modification in old mice was paralleled by downregulation of the glyoxalase network in all the investigated organs, which may, at least in part, explain the observed increase of CML-modified proteins in old mice. A similar age-associated decline of glyoxalases in rodents and humans has been reported earlier (Kuhla et al., 2006; Sharma-Luthra and Kale, 1994).

Our data reveal a substantial overlap between the identified CML sites and other PTM, particularly acetylation and ubiquitination, suggesting that CML modification may interfere with biological processes mediated by these PTM. For instance, K92 on histone H4, which became substantially more modified with CML during aging, has been reported to be also a target of acetylation and glutarylation, which are both involved in the regulation of chromatin structure and dynamics in response to DNA damage (Bao et al., 2019; Ye et al., 2005). Furthermore, global glycation of histone 3 has been shown to compete with acetylation and methylation thereby disrupting chromatin architecture (Zheng et al., 2019; Zheng et al., 2020). Thus, the age-dependent increase in CML modification of histone H4-K92 as seen in our study may contribute to alteration of chromatin and DNA damage responses in old tissues via interfering with other PTM.

In order to investigate other potential mechanisms by which AGEs might affect cellular phenotypes, we focused on perturbation of proteostasis. We show that glyoxal specifically affects the proteostasis network in MEF by increasing the abundance of UPS components and eliciting a hormetic effect on proteasome activity. Low doses of glyoxal increased the activity of the proteasome, while high doses did not alter proteasomal activity and led to an accumulation of ubiquitinated proteins. These data are in agreement with earlier reports showing that methylglyoxal is able to upregulate the protein quality control system (Zemva et al., 2017), and to impair the UPS at high levels leading to accumulation of toxic aggregates (Bento et al., 2010). In line with this, low doses of methylglyoxal led to an increased lifespan of *C. elegans*, while high doses (>1 mM) decreased lifespan (Ravichandran et al., 2018). Inhibition of proteasomal activity by dicarbonyls might be related to a direct modification of the 20S proteasome, as previously shown (Queisser et al., 2010). Here, we provide evidence for an interference of high AGEs with the stability of the proteasome by showing reduced thermal stability of proteasomal proteins in glyoxal-treated cells and identifying direct sites of CML modification of proteasomes both in cells treated with glyoxal as well as in organs obtained from mice. Interestingly, we also identified several enzymes involved in the ubiquitin cycle, including ubiquitin itself, to be direct targets of CML modification, suggesting that the interference of AGEs might extend to other components of the UPS beyond the proteasome. Thus, given that the proteasome activity is known to decrease across tissues during aging (Friguet et al., 2000; Hipp et al., 2019; Kelmer Sacramento et al., 2020; Saez and Vilchez, 2014), increased CML levels during aging might contribute to proteostasis impairment via alteration of the UPS.

Finally, we investigated the impact of glyoxal on the functional phenotype of endothelial cells, which are well known to be involved in the pathogenesis of age-associated metabolic diseases (Gimbrone and Garcia-Cardena, 2016; Sena et al., 2013). In line with our initial proteomic analysis, we found that glyoxal triggered a proliferation inhibition phenotype characterized by reduced cell proliferation in the presence of serum or bFGF and by lower sprouting in response to VEGF. Glyoxal had only minor effects on endothelial energy metabolism without changes in cellular ATP levels and did not induce cytotoxic effects, as shown by the absence of apoptosis. Our data are in line with previous studies showing that dicarbonyls induce endothelial dysfunction via oxidative stress, inflammatory responses, senescence or apoptosis (Jang et al., 2017; Liu et al., 2012; Navarrete Santos et al., 2017; Sliman et al., 2010; Wang et al., 2019; Yamawaki and Hara, 2008) and that AGEs interfere via RAGE-dependent signaling processes in endothelial cells (Chen et al., 2020; Li et al., 2018; Ravi et al., 2020). Until now, however, the impact of dicarbonyl-induced changes of protein abundance and intracellular protein modifications on endothelial function has not been thoroughly investigated.

To unravel mechanisms underlying the glyoxal-induced proliferation inhibition phenotype, we looked at differentially expressed and/or CML-modified proteins related to cell cycle control. We found that various cell cycle regulators were downregulated in glyoxal-treated cells, particularly proteins involved in DNA replication, likely because CDKN1A/p21 was strongly upregulated and thus restricted S phase entry. This pattern was underlined by growth arrest in G0/G1 and G2 phases and by activation of a DNA damage response as possible underlying mechanism (Ciccia and Elledge, 2010; Shaltiel et al., 2015). We were able to detect phosphorylation of CHK1 as a marker for an activated G1 and S phase checkpoint and γH2AX foci as indicator of DNA damage, likely DNA double strand breaks (Kuo and Yang, 2008; Zhang and Hunter, 2014). This was paralleled by an upregulation of the p53/p21 pathway, a major DNA damage response effector known to prevent G1/S transition (Abbas and Dutta, 2009; Helton and Chen, 2007). Glyoxal may cause DNA damage by glycation of DNA leading to so-called nucleotide AGEs (Pischetsrieder et al., 1999; Rabbani and Thornalley, 2015; Thornalley, 2008; Waris et al., 2015) or by triggering the formation of ROS (Ahmad et al., 2018). The latter may have contributed to DNA damage in our study, since both mitochondrial and cytosolic ROS levels were enhanced by glyoxal. In addition to the p53/p21 pathway, the enhanced expression of p16, another inhibitor of cyclin-dependent kinases preventing G1/S transition, is likely to contribute to the observed sustained growth arrest (Rayess et al., 2012). Upregulation of p16 may be linked to increased ROS levels induced by glyoxal (Sasaki et al., 2014; Takahashi et al., 2006). Both, p53/p21 and p16 pathways are known to play a role in the induction of premature senescence dependent on the extent and persistence of stress (Campisi and d’Adda di Fagagna, 2007) and may thus also be involved in mediating the moderate senescence response occurring in glyoxal-treated endothelial cells.

In addition to changes in protein abundance of cell cycle proteins induced by glyoxal, we observed CML modification of several proteins mainly involved in G2 and M phases of cell cycle. To understand how glycation of specific proteins may add to the glyoxal-induced inhibition of proliferation in endothelial cells, we focused on tubulin, whose α and β chains exhibited several CML modifications. Tubulin chains form protofilaments, which then associate to microtubules. The latter are major constituents of the cytoskeleton and form the mitotic spindle, whose function is to segregate chromosomes during cell division (Goodson and Jonasson, 2018). A major characteristic of tubulin polymers is their dynamic instability, i.e. the rapid transition between growth and shrinkage, which allows rapid reorganization of the cytoskeleton (Brouhard and Rice, 2018). Microtubule dynamics is known to be regulated by various PTM such as detyrosination, polyglutamination or acetylation (Song and Brady, 2015). Our data reveal glycation as a previously unknown tubulin modification, which has an impact on microtubule dynamics. We show that glyoxal treatment of endothelial cells triggered the formation of strongly structured tubulin filaments compared to fewer and thinner filaments in control cells, similar as observed with microtubule-stabilizing agents (Berges et al., 2017; Wang et al., 2017). Accordingly, our data reveal an increased stability of CML-modified microtubules. While the depolymerizing compound nocodazole completely dissolved the tubulin filaments in control cells, its effect was significantly reduced in cells pretreated with glyoxal. Microtubule dynamics may not only be affected by CML modification of tubulin but also by competition of glycation with other PTM. As shown in our proteomic analysis of mouse tissues, CML modification often occurs on residues that are known to be modified by other PTM. Accordingly, we found four CML-modified sites on α-tubulin (K326, K336, K394, K401) and three CML sites on β-tubulin (K58, K324, K379), which overlap with previously described acetylation sites (Sadoul and Khochbin, 2016). Moreover, we were able to demonstrate a direct competition between glycation and acetylation for K58 of β-tubulin indicating that exposure of endothelial cells to glyoxal may alter the tubulin code (Ferreira et al., 2018; Janke and Magiera, 2020). The latter refers to the concept that PTM of tubulin modulate the composition of individual microtubules and program them for specific functions, for instance by regulating the interaction with microtubule-binding proteins (Janke and Magiera, 2020). While the functional impact of most acetylation marks is not yet understood, acetylation of K40 within the microtubule lumen is suggested to protect microtubules from mechanical stress, possibly through affecting interaction between tubulin protofilaments (Portran et al., 2017; Xu et al., 2017). We found a decrease in K40 acetylation in glyoxal-treated cells, which adds to the alteration of the tubulin code although CML modification was not detected at this site. To address the question of whether the observed alteration of microtubule dynamics may affect mitosis, we compared mitosis in control and glyoxal-treated cells by employing a model, in which cells were treated with glyoxal after synchronization at the G1/S transition point. We found a lower expression of the mitotic marker p-H3(S10) in the presence of glyoxal indicating inhibition of mitosis which may be attributable to impaired microtubule dynamics and contribute to the antiproliferative effect of glyoxal. Together, these data describe a novel potential mechanism of cell cycle inhibition via posttranslational CML modification of tubulin.

The described proliferation inhibition phenotype of endothelial cells induced by glyoxal is likely to be of pathophysiological relevance. It is well known that CML accumulates in tissues in patients with diabetes and/or cardiovascular diseases as well as in aging (Delgado-Andrade, 2016). On the other hand, reduced angiogenesis leading to impaired wound healing is one of the hallmarks of diabetes (Okonkwo and DiPietro, 2017) and also observed in aging (Hodges et al., 2018; Moriya and Minamino, 2017). It is also known that only fully functional endothelial cells are able to generate new blood vessels (Boodhwani and Sellke, 2009; Sun et al., 2009). Thus, the formation of glyoxal and its effect on endothelial cell proliferation may be one of the factors linking metabolic alterations to disturbed angiogenesis in diabetes and aging and promote pathogenetic processes via this mechanism.

Together, the data presented in this study are based on a novel enrichment strategy coupled to mass spectrometry, which allowed the identification of proteins susceptible to CML modification and their respective CML sites in cells treated with a glycating agent and in organs from mice. Our data unravel subsets of proteins prone to CML modification as well as a proteotoxic response characterized by a dysfunction of the UPS. Exposure of endothelial cells to glycative stress triggers a proliferation inhibition phenotype, in which altered expression of cell cycle regulators induced via the DNA damage response as well as CML modification of tubulin contribute to growth arrest. The observations of this study expand our understanding of vascular dysfunction in aging and age-related metabolic disease and pave the way for further studies to link CML modifications to functional phenotypes.

## Acknowledgments

The authors gratefully acknowledge support from the FLI Core Facilities Proteomics, FACS, functional genomics and the Mouse Facility. The authors acknowledge Helmut Pospiech and Johannes Jungwirth (FLI, Jena) for providing advice and tools for cell cycle analysis, Elke Teuscher (Institute of Molecular Cell Biology, Jena) for her excellent technical assistance and the isolation and culture of HUVEC, Claudia Ender and Amod Godbole (Institute of Molecular Cell Biology, Jena) for taking immunofluorescent pictures, Titus Lohfink (Institute of Chemistry-Food Chemistry, Halle) for CML analysis in HUVEC, Max Tiessen and Domenico Di Fraia (FLI Jena) for implementing the R shiny webserver, and Julia Heiby and Ellen Späth (FLI Jena) for proofreading the manuscript. RH receives funds from the Deutsche Forschungsgemeinschaft (DFG, RTG1715 and RTG2155). AO acknowledges funding from the DFG (RTG2155), the Else Kröner Fresenius Stiftung (award number: 2019_A79), the Deutsches Zentrum für Herz-Kreislaufforschung (award number: 81X2800193) and the Fritz-Thyssen foundation (award number: 10.20.1.022MN). This research was also supported by the European Regional Development Fund (Grant ID: EFRE HSB 2018 0019) and the federal state of Thuringia providing technical equipment. The FLI is a member of the Leibniz Association and is financially supported by the Federal Government of Germany and the State of Thuringia.

## Author contributions

Conceptualization: SDS, AO, RH. Investigation: SDS, KS, JMK, AL, TLR, TB, CM. Methodology: SDS, KS, JMK. Data analysis: SDS, KS, JMK, LP, TB, AO, AL, TLR, RH. Project administration: AO, RH. Resources: AO, RH, MAG. Supervision: ZQW, MAG, AO, RH. Visualization: SDS, KS, AO. Writing – original draft: SDS, KS, AO, RH. Writing – review & editing: TB, LP and ZQW.

## Material and Methods

### Chemicals

M199 was purchased from Lonza (Verviers, Belgium). Fetal calf serum (FCS), human serum, endothelial growth supplement (ECGS), glyoxal (HUVEC studies), hydroxyurea, nocodazole, antimycin A, thymidine, 2-deoxy-D-glucose, trypsin inhibitor, thrombin, aprotinin, 5-bromo-4-chloro-3-indolyl β-D-galactopyranoside (X-Gal), 4’,6-diamidino-2-phenylindole (DAPI), octyl β-D-glucopyranoside, iodoacetamide (IAA), aqueous NH_3_, ATP, cOmplete™, EDTA-free protease inhibitor cocktail, HEPES, MOPS, NP40 and PonceauS were purchased from Sigma (Taufkirchen, Germany). Protease inhibitor mixture complete, EDTA-free was obtained from Roche Diagnostics (Mannheim, Germany), fibrinogen from Merck/Millipore (Darmstadt, Germany) and Fluoromount-G^®^ from Southern Biotech (Birmingham, AL, US), respectively. Bovine serum albumin-C (BSA-C) was from Aurion (Wageningen, The Netherlands) and goat serum from Cell Signaling Technology (Frankfurt, Germany). D-glucose, L-glutamine, sodium pyruvate, 5-ethynyl-2’-deoxyuridine, Trypsin-EDTA and anhydrous dimethylsulfoxide (DMSO) derived from Thermo Fisher Scientific (Waltham, MA, USA). Oligomycin and carbonyl cyanide-4- (trifluoromethoxy)phenylhydrazone (FCCP) came from Abcam (Cambridge, UK). Carboxyfluorescein succinimidyl ester (CFSE) was purchased from Invitrogen (Carlsbad, CA, US). Glyoxal (MEF studies), glycerine, β-mercaptoethanol, glucose, dithiothreitol (DTT), EDTA, formic acid, ammonium bicarbonate, Tris, bovine serum albumin, Tween-20 and Triton X-100 were obtained from Carl Roth GmbH (Karlsruhe, Germany). Trifluoroacetic acid, acetonitrile and 2-propanol were from Biosolve BV (Valkenswaard, The Netherlands) and glycine was from VWR International (Radnor, PA, USA).

### Antibodies

Antibodies raised against β-actin (HUVEC studies), cleaved caspase 3 (D175), caspase 3, cleaved PARP (D214), PARP, histone 3, p-Chk1 (S317), p-Chk1 (S345), Chk1, γH2A.X (S139), p-p53 (S15), p53, p21, p16, α/β-tubulin and ac-tubulin (K40) were obtained from Cell Signaling Technology (Frankfurt, Germany). Antibodies against CML (western blot) and p-H3 (S10) (western blot) were from Abcam (Cambridge, UK). The antibody against p-H3 (S10) for flow cytometry was obtained from Merck/Millipore (Darmstadt, Germany) and cyclin A antibody was from Santa Cruz Biotechnology (Dallas, TX, US). The CML antibody used for enrichment was purchased from ImmuneChem Pharmaceuticals Inc. (Burnaby, Canada). Antibodies against mono- and K29-, K48-, and K63-linked mono- and polyubiquitinylated proteins were from Enzo Life Sciences (Farmingdale, NY, USA) and the antibody against β-actin (MEF studies) was from Sigma (Taufkirchen, Germany). Peroxidase-labeled anti-mouse and anti-rabbit IgG were from Kirkegaard and Perry Laboratories, Inc. (Gaithersburg, MD, USA) or from Dako GmbH (Hamburg, Germany). AlexaFluor^®^647-conjugated azide (AF647 azide) and secondary AlexaFluor^®^488-conjugated goat anti-rabbit IgG were from Thermo Scientific (Waltham, MA, USA).

### Mice

All wild-type mice were C57BL/6J obtained from Janvier Labs (Le Genest-Saint-Isle, France) or from internal breeding at FLI. All animals were kept in a specific pathogen-free animal facility with a 12 h light/dark cycle. Young mice were aged 3-4 months, old mice were aged 26-33 months. Mice had unlimited access to food (ssniff, Soest, Germany)) during the experiment. For the analysis of total proteome, CMLpepIP and glyoxalase activity measurements only male mice were used. Mice were euthanized and organs were isolated, washed in PBS, weighted and immediately snap-frozen in liquid nitrogen before storage in −80°C.

### MEF cell culturing and treatments

SV40-immortalized mouse embryonic fibroblasts (MEF, a kind gift of K. L. Rudolph) were grown in high glucose DMEM medium (Sigma, D6429) until they reached 70 % confluency. For glyoxal treatment, DMEM medium was removed, cells were washed with PBS and live cell imaging medium (140 mM NaCl; 2.5 mM KCl; 1.8 mM CaCl_2_; 1.0 mM MgCl_2_), 20 mM HEPES; 4500 mg/l glucose and pH 7.4) was added. Subsequently, glyoxal diluted in the same medium was added at final concentrations of 0.5 or 2 mM for 8 - 24 h, as notified in figures. For all treatments, cells were snap-frozen in liquid nitrogen and stored at −80°C until further processing.

### HUVEC cell culture and treatment

HUVEC were isolated from anonymously acquired human umbilical cords according to the “Ethical principles for Medical Research Involving Human Subjects” (Declaration of Helsinki 1964) as previously described (Spengler et al., 2020). The protocol was approved by the Jena University Hospital Ethics Committee. The donors were informed and gave written consent. Briefly, after rinsing the cord veins with 0.9 % NaCl, endothelial cells were detached with collagenase (0.01 %, 3 min at 37 °C), suspended in M199/10 % FCS, washed once (500 x g, 6 min) and seeded on a cell culture flask coated with 0.2 % gelatin. Full growth medium (M199, 17.5 % FCS, 2.5 % human serum, 7.5 μg/ml ECGS, 7.5 U/ml heparin, 680 μM glutamine, 100 μM vitamin C, 100 U/ml penicillin, 100 μg/ml streptomycin) was added 24 h later. HUVEC from the second passage were detached with trypsin/EDTA after two washes in PBS and the reaction was stopped with HEPES buffer (10 mM HEPES (pH 7.4), 145 mM NaCl, 5 mM KCl, 1 mM MgSO_4_, 1.5 mM CaCl_2_, 10 mM glucose) containing 10 % FCS (HEPES/FCS). Cells were counted using a Neubauer chamber. Seeding was carried out at densities between 23,000/cm^2^ and 27,500/cm^2^ dependent on whether experiments were performed 48 or 72 h after seeding, respectively. For senescence and growth experiments, the seeding density was lower (8,000/cm2 (senescence-associated β-galactosidase (SA-β-Gal) staining), 10,000/cm2 (BrdU incorporation), 15,000/cm2 (cell cycle), 20,500/cm2 (thymidine block)). If not otherwise indicated, 30 mm-dishes were used for experiments. For mass spectrometry-based proteomics, detachment of HUVEC from 90 mm-dishes was performed with trypsin/EDTA and the reaction was stopped by an equal amount of trypsin inhibitor (1 mg/ml in PBS). Then, cells were washed twice in M199 (500 x g, 3 min) and snap-frozen in liquid nitrogen. Treatment of HUVEC with glyoxal was performed in experimental medium (i.e. full growth medium without vitamin C) and started 48 - 72 h after seeding. Incubations were performed with glyoxal concentrations of 0.5 or 1 mM for 4 - 48 h, as notified in figures. Thereafter, cells were stimulated with growth factors if indicated and processed as described below.

### Sample preparation for quantitation of total CML levels

MEF and mouse organs were lysed and proteins acetone precipitated as described in *Sample preparation for mass spectrometry-based proteomics*. HUVEC were detached by trypsin/EDTA, transferred to HEPES/FCS, centrifuged (500 x *g*, 6 min), washed twice in 2 ml PBS and subjected to one freezing/thawing cycle in liquid nitrogen. Pellets were resuspended in 200 μl ice-cold Tris buffer (50 mM Tris (pH 7.4), 2 mM EDTA, 1 mM EGTA, 50 mM NaF, 150 mM NaCl, 10 mM Na_4_P_2_O_7_, 1 mM Na_3_VO_4_, 1 % Triton X-100, 0.1 % SDS, 0.5 % sodium deoxycholate, 1 mM PMSF, 10 μl/ml protease inhibitor cocktail) and incubated on ice for 30 min. After homogenization using a tissue homogenizer, proteins were precipitated by adding TCA to a final concentration of 10 %. Samples were centrifuged (4000 x *g*, 5 min, 4 °C), pellets were washed twice in ice-cold 80 % acetone and stored at −80 °C until further processing.

### Quantitation of total CML levels by HPLC-MS/MS

Protein pellets were reconstituted in PBS and 250 μl aliquots of protein extracts (1 mg/ml) were reduced by addition of 100 μl NaBH_4_ solution (15 mg/ml in 0.01 M NaOH) and were shaken for 1 h at room temperature. Samples were dried in a vacuum concentrator (Savant-Speed-Vac Plus SC 110 A combined with a Vapor Trap RVT 400, Thermo Fisher Scientific, Bremen, Germany). 800 μl of 6 M HCl was added and the solution was heated 20 h at 110°C under an argon atmosphere. Volatiles were removed in a vacuum concentrator and the residue was dissolved in 300 μl of ultra-pure water. Samples were filtered through 0.45 μm cellulose acetate Costar SpinX filters (Corning Inc., Corning, USA). After complete hydrolysis, the amount of amino acids in hydrolysates was determined by ninhydrin assay and referenced to a calibration of l-leucine concentrated between 5 and 100 μM as described previously (Smuda et al., 2015). The absorbance was determined at 546 nm with an Infinite M200 microplate reader (Tecan, Männedorf, Switzerland) using 96-well plates.

Chromatographic separations were performed on a stainless-steel column (XSelect HSS T3, 250×3.0mm, RP18, 5μm, Waters, Milford, USA) using a flow rate of 0.7 ml/min and a column temperature of 25 °C. Eluents were ultra-pure water (A) and a mixture of methanol (Biosolve, 0013684102BS) and ultra-pure water (7:3, (v/v); B), both supplemented with 1.2 ml/l heptafluorobutyric acid. Samples were injected (10 μl) at 2% B and run isocratic for 2 min, gradient was changed to 14 % B within 10 min (held for 0 min), 87 % B within 22 min (held for 0 min), 100% B within 0.5 min (held for 7 min) and 2 % B within 2.5 min (held 8 min).

A PU-2080 Plus quaternary gradient pump with degasser and an AS-2057 Plus autosampler (Jasco, Gross-Umstadt, Germany) were used. The mass analyses were performed using an API 4000 quadrupole instrument (Applied Biosystems, Foster City, USA) equipped with an API source using electrospray ionization. The HPLC system was connected directly to the probe of the mass spectrometer. Nitrogen was used as sheath and auxiliary gas. To measure CML the scheduled multiple-reaction monitoring (sMRM) mode of HPLC-MS/MS was used. Quantitation was based on the standard addition method using known amounts of pure CML standard to compensate for matrix effects. The authentic reference compound was added at 0.5, 1, 2, and 4 times the concentration of the analyte in the sample and correlation coefficients were 0.9 or higher.

### Thermal Proteome Profiling (TPP)

MEF were grown in 150 mm-dishes until they reached 70-80 % confluence. They were treated at different concentration of glyoxal (0, 0.5 mM and 2 mM) for 8 h in imaging medium (as described in “*MEF cells culturing and treatments*”). Cells were harvested using Trypsin-EDTA phenol red (0.05 %) and counted in order to have 10^6^ cells per condition in PBS. Thereupon, cell suspensions were split into 10 individual 0.2 ml PCR tubes in equal volumes (100 μl each tube) and quickly span down to reach a final volume of 20 μl of cell suspension in each tube. Tubes were heated at different temperatures for 3 min using a 96 well dry bath ThermoQ (Hangzhou Bioer Technology, Hangzhou, China). Temperatures were: 37, 41, 44, 47, 50, 53, 56, 60, 63 and 67 °C. Cells were then incubated at 25 °C for 3 min, before adding 80 μl of 0.625 % NP40 and being snap-frozen in liquid nitrogen. Cells were lysed by a thaw-freeze-thaw cycle composed of an incubation step for 5 min at 25 °C, followed by snap freezing, and an additional incubation at 25 °C for 5 min. Lysates were transferred to 7 x 20mm polycarbonate thick-wall tubes (Beckman Coulter, Krefeld, Germany) for ultra-centrifugation (centrifuge Optima TLX with rotor TLA 100, Beckman Coulter), which was carried out at 100,000 x *g* for 20 min at 4 °C. Identical volumes of the resulting supernatants were then taken (32 μl) estimated to correspond to approximately 30 μg of protein extract for the 37 °C sample. Samples were reduced with 10 mM DTT for 15 min at 45°C and cysteine alkylated with freshly prepared 15 mM IAA for 30min at 25 °C, in the dark. Following reduction and alkylation, proteins were precipitated with ice-cold acetone and digested into peptides, as described in “*Sample preparation for mass spectrometry-based proteomics*”.

### Sample preparation for mass spectrometry-based proteomics

HUVEC and MEF cell pellets (300k and 100k respectively) were thawed, reconstituted in 50 μl of ice-cold PBS and lysed by addition of 50 μl 2 x lysis buffer (2% SDS, 100 mM HEPES pH 8, 20 mM DTT). Mouse organs were thawed and transferred into Precellys^®^ lysing kit tubes (Keramik-kit 1.4/2.8 mm, 2 ml (CKM)) containing 1 ml of PBS supplemented with 1 tab of cOmplete™, Mini, EDTA-free Protease Inhibitor per 50 ml. For homogenization, tissues were shaken twice at 6000 rpm for 30 s using Precellys^®^ 24 Dual (Bertin Instruments, Montigny-le-Bretonneux, France) and the homogenate was transferred to new 2 ml Eppendorf tubes. Based on estimated protein content (5 % of fresh tissue weight for liver and 8 % for heart and kidney), 100 μg of protein was processed for further analyses. Volumes were adjusted using PBS and one volume equivalent of 2x lysis buffer was added.

Samples were sonicated in a Bioruptor Plus (Diagenode, Seraing, Belgium) for 10 cycles with 1 min ON and 30 s OFF with high intensity at 20 °C. Samples were quickly centrifuged and a second sonication cycle was performed as described above. The lysates were centrifuged at 18,407 x *g* for 1 min and transferred to new 1.5 ml Eppendorf tubes. Subsequently, samples were reduced using 10 mM DTT for 30 min at room temperature, and alkylated using freshly made 15 mM IAA for 30 min at room temperature in the dark. Subsequently, proteins were acetone precipitated and digested using LysC (Wako sequencing grade) and trypsin (Promega sequencing grade), as described in (Buczak et al., 2020). The digested proteins were then acidified with 10 % (v/v) trifluoracetic acid and desalted using *Waters Oasis^®^ HLB μElution Plate 30 μm* following manufacturer instructions. The eluates were dried down using a vacuum concentrator, and reconstituted samples in 5 % (v/v) acetonitrile, 0.1 % (v/v) formic acid. For Data Independent Acquisition (DIA) based analysis, samples were transferred to an MS vial, diluted to a concentration of 1 μg/μl, and spiked with iRT kit peptides (Biognosys, Zurich, Switzerland) prior to analysis by LC-MS/MS. For Tandem Mass Tags (TMT) based analysis, samples were further processed for TMT labelling as described below.

### TMT labelling and high pH peptide fractionation for organ aging proteome and TPP

Reconstituted peptides (at 1 μg/μl) were buffered using 100 mM HEPES buffer pH 8.5 (1:1 ratio) for labelling. 10-20 μg peptides were taken for each labelling reaction. TMT-10plex reagents (Thermo Scientific, Waltham, MA, USA) were reconstituted in 41 μl 100 % anhydrous DMSO. TMT labeling was performed by addition of 1.5 μl of the TMT reagent. After 30 min of incubation at room temperature with shaking at 600 rpm in a thermomixer (Eppendorf, Hamburg, Germany), a second portion of TMT reagent (1.5 μl) was added and incubated for another 30 min. After checking labelling efficiency, samples were pooled (45-50 μg total), desalted with *Oasis^®^ HLB μElution Plate* and subjected to high pH fractionation prior to MS analysis.

Offline high pH reverse phase fractionation was performed using a Waters XBridge C18 column (3.5 μm, 100 x 1.0 mm, Waters) with a Gemini C18, 4 x 2.0 mm SecurityGuard (Phenomenex) cartridge as a guard column on an Agilent 1260 Infinity HPLC, as described in (Buczak et al., 2020). Forty-eight fractions were collected along with the LC separation, which were subsequently pooled into 16 fractions (for liver and kidney) and 24 fractions (heart). Pooled fractions were dried in a vacuum concentrator and then stored at −80°C until LC-MS/MS analysis.

### LC-MS/MS based on Data Independent Acquisition (DIA) for MEF and HUVEC

Peptides (approx. 1 μg) were separated using a nanoAcquity UPLC M-Class system (Waters Milford, USA) with a trapping (nanoAcquity Symmetry C18, 5 μm, 180 μm x 20 mm) and an analytical column (nanoAcquity BEH C18, 1.7 μm, 75 μm x 250 mm). The outlet of the analytical column was coupled directly to a Q-exactive HF-X (Thermo Fisher, Waltham, MA, USA) using the Proxeon nanospray source. Solvent A was water, 0.1% formic acid and solvent B was acetonitrile, 0.1 % formic acid. The samples (approx. 1 μg) were loaded onto the trapping column with a constant flow of solvent A at 5 μl/min. Trapping time was 6 min. Peptides were eluted via the analytical column with a constant flow of 0.3 μl/min. During the elution step, the percentage of solvent B increased in a non-linear fashion from 0 % to 40 % in 90 min. Total runtime was 115 min, including clean-up and column re-equilibration. The peptides were introduced into the MS via a Pico-Tip Emitter 360 μm OD x 20 μm ID; 10 μm tip and a spray voltage of 2.2 kV was applied. The capillary temperature was set at 300 °C. The radio frequency (RF) ion funnel was set to 40 %.

Data from a subset of conditions were first acquired in data-dependent acquisition (DDA) mode to contribute to a sample specific spectral library. The conditions were as follows: Full scan MS spectra with mass range 350-1650 m/z were acquired in profile mode in the Orbitrap with resolution of 60,000 FWHM. The filling time was set at maximum of 20 ms with limitation of 3 x 10^6^ ions. The “Top N” method was employed to take the 15 most intense precursor ions (with an intensity threshold of 4 x 10^4^) from the full scan MS for fragmentation (using HCD normalized collision energy, 27 %) and quadrupole isolation (1.6 Da window) and measurement in the Orbitrap (resolution 15,000 FWHM, fixed first mass 120 m/z). The peptide match ‘preferred’ option was selected and the fragmentation was performed after accumulation of 2 x 10^5^ ions or after filling time of 25 ms for each precursor ion (whichever occurred first). MS/MS data were acquired in profile mode. Only multiply charged (2+ - 5+) precursor ions were selected for MS/MS. Dynamic exclusion was employed with maximum retention period of 20 s and relative mass window of 10 ppm. Isotopes were excluded. In order to improve the mass accuracy, internal lock mass correction using a background ion (m/z 445.12003) was applied.

For DIA, the same gradient conditions were applied to the LC as for the DDA and the MS conditions were varied as described: Full scan MS spectra with mass range 350-1650 m/z were acquired in profile mode in the Orbitrap with resolution of 120,000 FWHM. The default charge state was set to 3+. The filling time was set at maximum of 60 ms with limitation of 3 x 10^6^ ions. DIA scans were acquired with 34 mass window segments of differing widths across the MS1 mass range. HCD fragmentation (stepped normalized collision energy; 25.5, 27, 30 %) was applied and MS/MS spectra were acquired with a resolution of 30,000 FWHM with a fixed first mass of 200 m/z after accumulation of 3 x 10^6^ ions or after filling time of 40 ms (whichever occurred first). Data were acquired in profile mode. For data acquisition and processing of the raw data Xcalibur 4.0 and Tune version 2.9 (both Thermo Fisher) were employed.

### Data processing for DIA

DpD (DDA plus DIA) libraries were then created by searching both the DDA runs and the DIA runs using Spectronaut Pulsar (v10/11, Biognosys, Zurich, Switzerland). The data were searched against species specific protein databases (Uniprot *Mus musculus* or *Homo sapiens*, reviewed entry only, release 2016_01) with a list of common contaminants appended. The data were searched with the following modifications: carbamidomethyl (C) as fixed modification, and oxidation (M), acetyl (protein N-term) and carboxymethyllysine (CML) as variable modifications. A maximum of 3 missed cleavages was allowed. The library search was set to 1 % false discovery rate (FDR) at both protein and peptide levels. This library contained 92,917 precursors, corresponding to 5,393 protein groups for HUVEC, and 83,207 precursors, corresponding to 4,725 protein groups for MEF using Spectronaut protein inference. DIA data were then uploaded and searched against this spectral library using Spectronaut Professional (v.11) and default settings. Relative quantification was performed in Spectronaut for each pairwise comparison using the replicate samples from each condition using default settings, except: Major Group Quantity = Sum peptide quantity; Major Group Top N = OFF; Minor Group Quantity = Sum precursor quantity; Minor Group Top N = OFF; Data Filtering = Q value sparse; Normalization Strategy = Local normalization; Row Selection = Q value complete. The data (candidate tables) and protein quantity data reports were then exported and further data analyses and visualization were performed with R (v.3.6.3) and R studio server (v. 1.2.5042) using in-house pipelines and scripts.

### LC-MS/MS based on Tandem Mass Tags (TMT) for organ aging proteome and TPP

For TMT experiments, fractions were resuspended in 10 μl reconstitution buffer (5% (v/v) acetonitrile, 0.1 % (v/v) trifluoroacetic acid in water) and 3 μl were injected. Peptides were analyzed as described in (Buczak et al., 2020) using a nanoAcquity UPLC system (Waters) fitted with a trapping (nanoAcquity Symmetry C18, 5 μm, 180 μm x 20 mm) and an analytical column (nanoAcquity BEH C18, 2.5 μm, 75 μm x 250 mm), and coupled to an Orbitrap Fusion Lumos (Thermo Fisher Scientific, Waltham, MA, USA). Briefly, peptides were separated using a 130 min nonlinear gradient. Full scan MS spectra with mass range 375-1500 m/z were acquired in profile mode in the Orbitrap with resolution of 60,000 FWHM using the quad isolation. The RF on the ion funnel was set to 40 %. The filling time was set at maximum of 100 ms with an AGC target of 4 x 10^5^ ions and 1 microscan. The peptide monoisotopic precursor selection was enabled along with relaxed restrictions if too few precursors were found. The most intense ions (instrument operated for a 3 s cycle time) from the full scan MS were selected for MS2, using quadrupole isolation and a window of 1Da. HCD was performed with collision energy of 35 %. A maximum fill time of 50 ms for each precursor ion was set. MS2 data were acquired with fixed first mass of 120 m/z. The dynamic exclusion list was with a maximum retention period of 60 sec and relative mass window of 10 ppm. The instrument was not set to inject ions for all available parallelizable time. For the MS3, the precursor selection window was set to the range 400-2000 m/z, with an exclude width of 18 m/z (high) and 5 m/z (low). The most intense fragments from the MS2 experiment were co-isolated (using Synchronus Precursor Selection=8) and fragmented using HCD (65 %). MS3 spectra were acquired in the Orbitrap over the mass range 100-1000 m/z and resolution set to 30000. The maximum injection time was set to 105 ms and the instrument was set not to inject ions for all available parallelizable time.

### Data processing for TMT

TMT-10plex data from aging mouse organs were processed using Proteome Discoverer v2.0 (Thermo Fisher Scientific, Waltham, MA, USA). Data were searched against the fasta database (Uniprot *Mus musculus* database, reviewed entry only, release 2016_01) using Mascot v2.5.1 (Matrix Science) with the following settings: Enzyme was set to trypsin, with up to 1 missed cleavage. MS1 mass tolerance was set to 10ppm and MS2 to 0.5Da. Carbamidomethyl cysteine was set as a fixed modification and oxidation of methionine as variable. Other modifications included the TMT-10plex modification from the quantification method used. The quantification method was set for reporter ions quantification with HCD and MS3 (mass tolerance, 20 ppm). The false discovery rate for peptide-spectrum matches (PSMs) was set to 0.01 using Percolator (Brosch et al., 2009).

Reporter ion intensity values for the PSMs were exported and processed with procedures written in R (v.3.6.3) and R studio server (v. 1.2.5042), as described in (Heinze et al., 2018). Briefly, PSMs mapping to reverse or contaminant hits, or having a Mascot score below 15, or having reporter ion intensities below 1 x 10^3^ in all the relevant TMT channels were discarded. TMT channels intensities from the retained PSMs were then log2 transformed, normalized and summarized into protein group quantities by taking the median value using MSnbase (Gatto and Lilley, 2012). At least two unique peptides per protein were required for the identification and only those peptides with no missing values across all 10 channels were considered for quantification. Protein differential expression was evaluated using the limma package (Ritchie et al., 2015). Differences in protein abundances were statistically determined using the Student’s t test moderated by the empirical Bayes method. P values were adjusted for multiple testing using the Benjamini-Hochberg method (FDR, denoted as “adj. p”) (Benjamini and Hochberg, 1995).

### Data processing for TPP

For the analysis of the TPP sample data (TMT10plex, high pH fractionated, MS3 data acquisition), raw data files were first processed through the preMascot process of the isobarquant package (Franken et al., 2015). After the .mgf files had been generated, these were processed via Mascot Daemon (Matrix Science) using Mascot version 2.5.1. Firstly, species-specific database including decoy was created by concatenating the forward entries (Uniprot *Mus musculus* database, reviewed entry only, release 2016_01) to the reversed sequences of the database. Data were then searched against the relevant database with the following settings: Enzyme = trypsin, 3 missed cleavage allowed. Modifications: carbamidomethyl (C) and TMT10plex (K) as fixed modifications; oxidation (M), carboxymethyllysine (K) and TMT10plex (N-term) as variable. MS1 tolerance was 10 ppm, MS2; 0.5 Da. When the .dat files for each fraction had been generated, these were subjected to the postMascot process of isobarquant, combining all the outputs into a single, merged, TMT10plex quantified protein output for further processing of melting point curves using the TPP package in R (Childs et al., 2019).

### Enrichment of CML modified peptides (CMLpepIP)

Lysates containing approximatively 1 mg protein for HUVEC and MEF and 5 mg for tissues were digested as described in “*Sample preparation for mass spectrometry-based proteomics*” with minor modifications: the digestion buffer used was 3 M Urea, 100 mM HEPES, 5 % (v/v) acetonitrile and the ratio of LysC and Trypsin used was 1:150 enzyme to protein. The digests were then acidified with 10 % (v/v) trifluoroacetic acid and then desalted with *Waters Oasis^®^ HLB 96-well Plate 30 μm* (Waters Corp., Milford, MA, USA) in the presence of a slow vacuum. In this process, the columns were conditioned with 2x 1000 μl solvent B (80% (v/v) acetonitrile; 0.05 % (v/v) formic acid) and equilibrated with 2x 1000 μl solvent A (0.05% (v/v) formic acid in milliQ water). The samples were loaded, washed 2 times with 1000 μl solvent A, and then eluted with 500 μl solvent B. The eluates were dried down and dissolved in 200 μl of IP buffer (50 mM MOPS, pH 7.3, 10 mM KPO_4_ pH 7.5, 50 mM HEPES, 2.5 mM octyl β-D-glucopyranoside) at concentration of 5 μg/μl followed by sonication in a Bioruptor Plus (5 cycles with 1 min ON and 30 s OFF with high intensity at 20 °C). 10 % of the sample was kept and analyzed separately as input control. A pre-cleaning step was applied by incubating each sample with 10 μl of Protein A magnetic beads (New England Biolabs GmbH, Frankfurt, Germany) for 1 h at 4 °C in tube roller (15 rpm) (STARLAB Tube roller Mixer RM Multi-1). Samples were transferred to a magnetic rack (DynaMag™-2, Invitrogen). The supernatant was transferred into a new 1.5 ml Eppendorf tube and incubated overnight with 30 μg of pan (ε-N) CML antibody at 4 °C on tube roller (15 rpm) (STARLAB Tube roller Mixer RM Multi-1). Subsequently, 150 μl of Protein A magnetic beads were added to the samples and incubated for 1 h at 4°C on tube roller as previously. Samples were then transferred to a magnetic rack and the flow through was collected in a fresh tube, and beads were washed 3 times in 300 μl IP buffer. The enriched peptides were then eluted 3 times in 54 μl of 0.1 M glycine pH 2.6. The fraction of elution, flow through and the input were desalted using Macro Spin Column C18 columns (Harvard Apparatus, Cambridge, MA, USA) following manufacturer instructions. The eluates were then dried down in a vacuum concentrator, dissolved in in 5 % (v/v) acetonitrile, 0.1 % (v/v) formic acid and directly analyzed by LC-MS/MS for the MEF and HUVEC. An additional step of high pH peptide fractionation was performed for eluates from mouse organs, as described in “*TMT labelling and high pH peptide fractionation for organ aging proteome and TPP*”. Eight pooled fractions from high pH fractionation were measured by LC-MS/MS for each sample.

### LC-MS/MS based on Data Dependent Acquisition (DDA) for CMLpepIP

Peptides were separated using the nanoAcquity UPLC system (Waters) fitted with a trapping (nanoAcquity Symmetry C18, 5 μm, 180 μm x 20 mm) and an analytical column (nanoAcquity BEH C18, 1.7 μm, 75 μm x 250 mm). The outlet of the analytical column was coupled directly to an Orbitrap Fusion Lumos (Thermo Fisher Scientific, Waltham, MA, USA) using the Proxeon nanospray source. Solvent A was water, 0.1 % (v/v) formic acid and solvent B was acetonitrile, 0.1 % (v/v) formic acid. The samples (500 ng) were loaded with a constant flow of solvent A at 5 μl/min onto the trapping column. Trapping time was 6 min. Peptides were eluted via the analytical column with a constant flow of 0.3 μl/min. During the elution step, the percentage of solvent B increased in a linear fashion from 3 % to 25 % in 30 min, then increased to 32 % in 5 more minutes and finally to 50 % in a further 0.1 min. Total runtime was 60 min. The peptides were introduced into the mass spectrometer via a Pico-Tip Emitter 360 μm OD x 20 μm ID; 10 μm tip (New Objective) and a spray voltage of 2.2 kV was applied. The capillary temperature was set at 300 °C. The RF lens was set to 30 %. Full scan MS spectra with mass range 375-1500 m/z were acquired in profile mode in the Orbitrap with resolution of 120,000 FWHM. The filling time was set at maximum of 50 ms with limitation of 2 x 10^5^ ions. The “Top Speed” method was employed to take the maximum number of precursor ions (with an intensity threshold of 5 x 10^3^) from the full scan MS for fragmentation (using HCD collision energy, 30 %) and quadrupole isolation (1.4 Da window) and measurement in the ion trap, with a cycle time of 3 s. The MIPS (monoisotopic precursor selection) peptide algorithm was employed but with relaxed restrictions when too few precursors meeting the criteria were found. The fragmentation was performed after accumulation of 2 x 10^3^ ions or after filling time of 300 ms for each precursor ion (whichever occurred first). MS/MS data were acquired in centroid mode, with the rapid scan rate and a fixed first mass of 120 m/z. Only multiply charged (2+ - 7+) precursor ions were selected for MS/MS. Dynamic exclusion was employed with maximum retention period of 60 s and relative mass window of 10 ppm. Isotopes were excluded. Additionally, only 1 data dependent scan was performed per precursor (only the most intense charge state selected). Ions were injected for all available parallelizable time. In order to improve the mass accuracy, a lock mass correction using a background ion (m/z 445.12003) was applied. For data acquisition and processing of the raw data, Xcalibur 4.0 (Thermo Scientific, Waltham, MS, USA) and Tune version 2.1 were employed.

### Data processing for DDA for CMLpepIP

Software MaxQuant (version 1.5.3.28) was used to search the data. The data were searched against species-specific databases (Uniprot *Mus musculus* or *Homo sapiens*, reviewed entry only, release 2016_01) with a list of common contaminants appended. The data were searched with the following modifications: carbamidomethyl (C) as fixed modification, and oxidation (M), acetyl (protein N-term) and carboxymethyllysine (CML) as variable modifications. The mass error tolerance for the full scan MS spectra was set at 20 ppm and for the MS/MS spectra at 0.5 Da. A maximum of 3 missed cleavages were allowed. Peptide and protein level 1 % FDR were applied using a target-decoy strategy (Elias and Gygi, 2007). MaxQuant outputs were then used to perform qualitative analyses. In addition, highly confident CML sites were defined using these additional parameters: Posterior Error Probability <0.05, score >50, score difference >5, localization probability >0.75.

### Enrichment of acetylated peptides

HUVEC pellets (3 biological replicates per condition/time point) corresponding to approximately 500μg of proteins were used. Cell pellets were thawed, lysed and digested as described in the paragraph “*Sample preparation for mass spectrometry-based proteomics*”. The digested peptides were acidified with 10% (v/v) trifluoroacetic acid and then desalted with *Waters Oasis^®^ HLB 96-well Plate 30 μm* following manufacturer instructions. The eluates were dried down and dissolved in 500 μl of IP buffer (50 mM MOPS pH 7.3, 10 mM KPO_4_ pH 7.5, 50 mM NaCl, 2.5 mM Octyl β-D-glucopyranoside) to reach a peptide concentration of 1 μg/μl, followed by sonication in a Bioruptor (5 cycles with 1 min ON and 30 s OFF with high intensity at 20°C). 10 % of the sample was kept as it is in order to use it as input. Agarose beads coupled to antibody against acetyl lysine were washed three times with washing buffer (20 mM MOPS pH 7.4, 10 mM KPO_4_ pH 7.5, 50 mM NaCl) before incubation with each sample-peptide for 1.5 h on a rotating well at 750 rpm (STARLAB Tube roller Mixer RM Multi-1). Samples were transferred into Clearspin filter microtubes (0.22 μm) (Dominique Dutscher SAS, Brumath, France) and centrifuged at 4 °C for 1 min at 2,000 x *g*. Beads were washed first with IP buffer (three times), then with washing buffer (three times) and finally with 5 mM ammonium bicarbonate (three times). Thereupon, the enriched peptides were then eluted first in basic condition using 50 mM aqueous NH_3_, then using 0.1 % (v/v) trifluoroacetic acid in 10% (v/v) 2-propanol and finally with 0.1 % (v/v) trifluoroacetic acid. Elutions were then dried down and reconstituted in MS buffer A (5 % (v/v) acetonitrile, 0.1 % (v/v) formic acid), acidified with 10 % (v/v) trifluoroacetic acid and then desalted with *Waters Oasis^®^ HLB μElution Plate 30 μm*. Desalted peptides were finally dissolved in MS buffer A and analyze by LC-MS/MS.

### LC-MS/MS based on Data Independent Acquisition (DIA) for acetylated peptides

Peptides were separated using the nanoAcquity UPLC system (Waters) fitted with a trapping (nanoAcquity Symmetry C18, 5 μm, 180 μm x 20 mm) and an analytical column (nanoAcquity BEH C18, 1.7 μm, 75 μm x 250 mm). The outlet of the analytical column was coupled directly to an Orbitrap Fusion Lumos (Thermo Fisher Scientific, Waltham, MA. USA) using the Proxeon nanospray source. Solvent A was water, 0.1 % (v/v) formic acid and solvent B was acetonitrile, 0.1 % (v/v) formic acid. The samples (approx. 1 μg for DDA and 3 μg for DIA) were loaded with a constant flow of solvent A at 5 μl/min onto the trapping column. Trapping time was 6 min. Peptides were eluted via the analytical column with a constant flow of 0.3 μl/min. During the elution step, the percentage of solvent B increased in a non-linear fashion from 0 % to 40 % in 40 min. Total runtime was 60 min, including clean-up and column re-equilibration. The peptides were introduced into the mass spectrometer via a Pico-Tip Emitter 360 μm OD x 20 μm ID; 10 μm tip (New Objective) and a spray voltage of 2.2 kV was applied. The capillary temperature was set at 300 °C. The RF lens was set to 30 %. Data from each sample were first acquired in DDA mode. The conditions were as follows: Full scan MS spectra with mass range 350-1650 m/z were acquired in profile mode in the Orbitrap with resolution of 60,000 FWHM. The filling time was set at maximum of 50 ms with limitation of 2 x 10^5^ ions. The “Top Speed” method was employed to take the maximum number of precursor ions (with an intensity threshold of 5 x 10^4^) from the full scan MS for fragmentation (using HCD collision energy, 30 %) and quadrupole isolation (1.4 Da window) and measurement in the Orbitrap (resolution 15,000 FWHM, fixed first mass 120 m/z), with a cycle time of 3 s. The MIPS (monoisotopic precursor selection) peptide algorithm was employed but with relaxed restrictions when too few precursors meeting the criteria were found. The fragmentation was performed after accumulation of 2 x 10^5^ ions or after filling time of 22 ms for each precursor ion (whichever occurred first). MS/MS data were acquired in centroid mode. Only multiply charged (2^+^ - 7^+^) precursor ions were selected for MS/MS. Dynamic exclusion was employed with maximum retention period of 15 s and relative mass window of 10 ppm. Isotopes were excluded. In order to improve the mass accuracy, internal lock mass correction using a background ion (m/z 445.12003) was applied. For data acquisition and processing of the raw data Xcalibur 4.0 (Thermo Scientific, Waltham, MA, USA)) and Tune version 2.1 were employed.

For the DIA data acquisition the same gradient conditions were applied to the LC as for the DDA and the MS conditions were varied as described: Full scan MS spectra with mass range 350-1650m/z were acquired in profile mode in the Orbitrap with resolution of 120,000 FWHM. The filling time was set at maximum of 20 ms with limitation of 5 x 10^5^ ions. DIA scans were acquired with 30 mass window segments of differing widths across the MS1 mass range with a cycle time of 3 s. HCD fragmentation (30 % collision energy) was applied and MS/MS spectra were acquired in the Orbitrap with a resolution of 30,000 FWHM over the mass range 200-2000 m/z after accumulation of 2 x 10^5^ ions or after filling time of 70 ms (whichever occurred first). Ions were injected for all available parallelizable time). Data were acquired in profile mode.

### Data processing for DIA for acetylated peptides

DpD (DDA plus DIA) libraries were created by searching both DDA and DIA runs using Spectronaut Pulsar (v.13), as described in “*Data processing for DIA*“ with the following modification: acetyl (K) was included as variable modification. The library contained 17,051 precursors, corresponding to 3,227 protein groups using Spectronaut protein inference. DIA data were then uploaded and searched against this spectral library using Spectronaut Professional (v.13) and default settings. Intensities of precursors deriving from acetylated peptide were obtained from the peptide report table and filtered for a localization score >= 0.75 and further processed using in house written scripts in R (v.3.6.3) and R studio server (v. 1.2.5042). Intensities were summarized at the level of acetylation site by summing the intensities of all the precursors containing a given acetylation sites. Acetylation site intensities were normalized across runs by log2 transformation and median centering.

### Parallel Reaction Monitoring (PRM) for CML modified peptides

Twenty-four peptides containing CML-modification were selected among the most confident and consistently identified peptides from CMLpepIP and their isotopically labelled version (heavy Arginine (U-13C6;U-15N4) or Lysine (U-13C6; U-15N2) at the C-term was added) synthesized by JPT Peptide Technologies GmbH (Berlin, Germany). Peptides were delivered as lyophilized and reconstituted in 20 % (v/v) acetonitrile, 0.1 % (v/v) formic acid and further pooled together in a ratio 1:1. An aliquot of the pooled peptides, corresponding to approximatively 150 fmol per peptide, was analyzed by both DDA and DIA LC-MS/MS and used for assay generation using Spectrodive v.9 (Biognosys AG, Schlieren, Switzerland). Assays were successfully developed for 20 out of 22 targeted peptides.

Following assay development, the heavy synthetic peptide pool was used to evaluate the limit of detection (LOD) and limit of blank (LOB). Synthetic peptide pool was mixed with 200 ng/μl of digested yeast peptides at concentration from 45 fmol to 5760 fmol on column and measured in triplicate using PRM. Peptides were separated using a nanoAcquity UPLC M-Class system (Waters, Milfors, MA, USA)) with a trapping (nanoAcquity Symmetry C18, 5 μm, 180 μm x 20 mm) and an analytical column (nanoAcquity BEH C18, 1.7 μm, 75 μm x 250 mm). The outlet of the analytical column was coupled directly to a Q-exactive HF-X (Thermo Fisher, Waltham, MA, Germany) using the Proxeon nanospray source. Solvent A was water, 0.1 % (v/v) formic acid and solvent B was acetonitrile, 0.1 % (v/v) formic acid. Peptides were eluted via the analytical column with a constant flow of 0.3 μl/min. During the elution step, the percentage of solvent B increased in a non-linear fashion from 0 % to 40 % in 40 min. Total runtime was 60 min, including clean-up and column re-equilibration. PRM acquisition was performed in a scheduled fashion for the duration of the entire gradient (after instrument calibration in an unscheduled mode) using the ‘‘DIA’’ mode with the following settings: resolution 120,000 FWHM, AGC target 3 x 10^6^, maximum injection time (IT) 250 ms, isolation window 0.4 m/z. For each cycle, a ‘‘full MS’’ scan was acquired with the following settings: resolution 120,000 FWHM, AGC target 3 x 10^6^, maximum injection time (IT) 10 ms, scan range 350 to 1650 m/z. Peak group identification and quantification was performed using SpectroDive v9. Quantification was performed using spike-in approach. Thereby, the summed height of all the identified transitions was used to estimate the quantity of each peptide. LOQ and LOB was performed using the R package MSstat (Choi et al., 2014) using non-linear mode.

For CML site quantification in aging organs, not enriched digested peptides were spiked with synthetic heavy peptides at a concentration of 144 fmol/μl and analyzed by scheduled PRM as described above, using the following sub-panel:

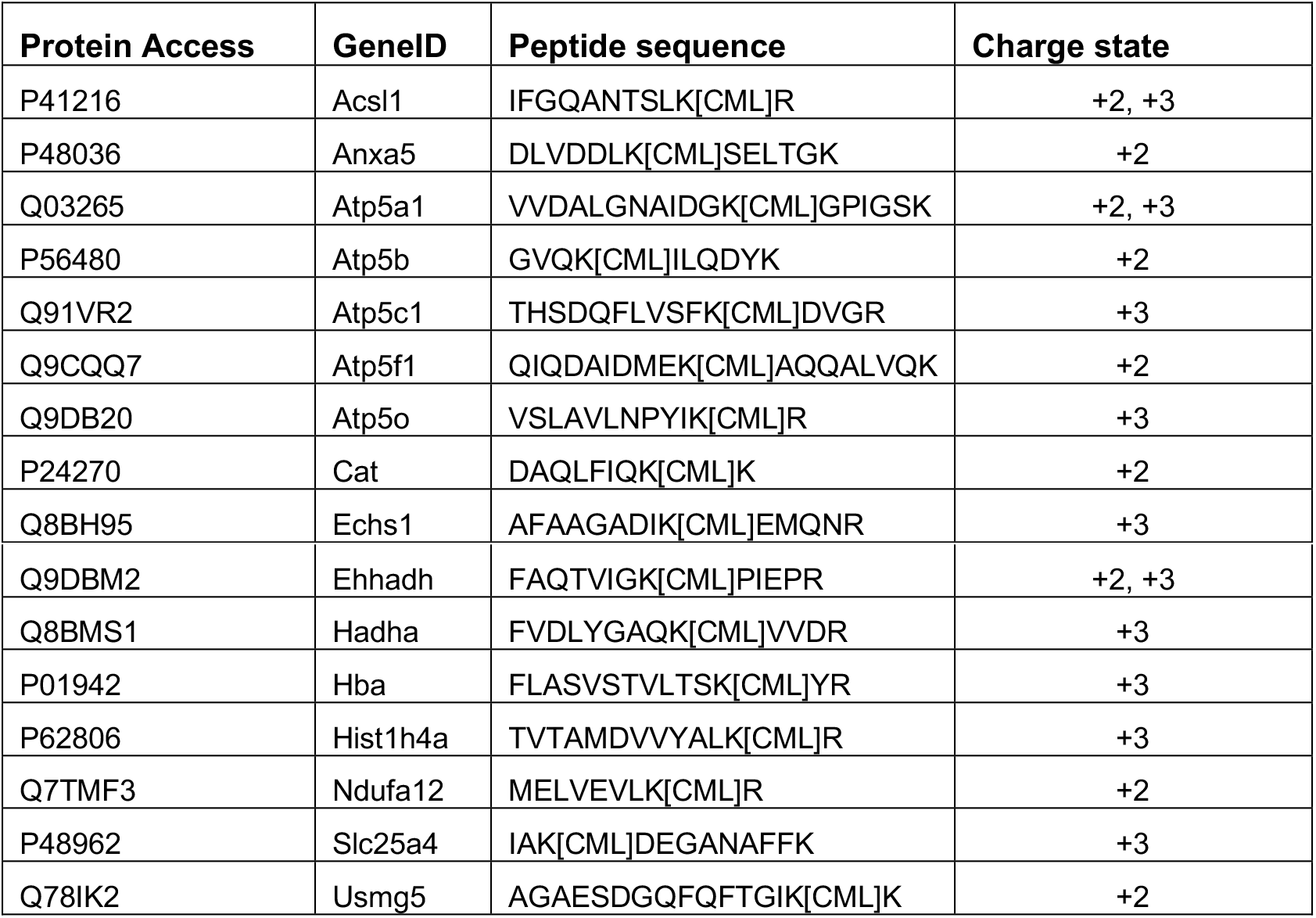

Peptides quantities were normalized across samples by dividing for the integrated intensity of the Base Peak Chromatogram extracted for each sample from “full MS” scans using Xcalibur v4.1.

### Immunoblot for mono and polyubiquitinated proteins of MEF cells treated with glyoxal

The same cell lysates used for proteasome activity assay were used. Lysates were thawed and centrifugated at 20817 x *g*, for 15 min at 4 °C to remove debris and the supernatant was transferred to a new tube. Based on EZQ assay performed on it, 20 μg of proteins were used. Samples were sonicated for 10 cycles (1min ON and 30s OFF) using a Bioruptor Plus with 4x loading buffer (1.5 M Tris .pH 6.8, 20% (w/v) SDS, 85 % (v/v) glycerin, 5 % (v/v) β-mercaptoethanol). Proteins were separated on 4–20 % Mini-Protean^®^ TGX™ Gels (BioRad, Neuberg, Germany) by sodium dodecyl sulfate polyacrylamide gel electrophoresis (SDS-PAGE) using a Mini-Protean^®^ Tetra Cell system (BioRad, Neuberg, Germany). Proteins were transferred to a nitrocellulose membrane (Millipore) using a Trans-Blot^®^ Turbo™ Transfer Starter System. Membranes were stained with PonceauS for 5 min on a shaker (Heidolph Duomax 1030), washed with milliQ water, imaged on a Molecular Imager ChemiDocTM XRS+ Imaging system (BioRad, Neuberg, Germany) and destained by 2 washes with PBS and 2 washes in TBST for 5 min (Tris-buffered saline (TBS, 25 mM Tris, 75 mM NaCl), with 0.5 % (v/v) Tween-20). After incubation for 1 h in blocking buffer (3 % bovine serum albumin (w/v) in TBST), membranes were stained overnight with primary antibodies against mono- and K29-, K48-, and K63-linked mono- and polyubiquitinylated proteins or β-actin diluted in blocking buffer (1:5000) at 4 °C on a tube roller (BioCote^®^ Stuart^®^ SRT6). Membranes were washed 3 times with TBST for 10 min at room temperature (RT) and incubated with horse radish peroxidase coupled secondary antibodies at RT for 1 h (1:2000 in 0.3 % (w/v) BSA in TBST). After 3 more washes for 10 min in TBST, chemiluminescent signals were detected using an ECL (enhanced chemiluminescence) detection kit (Thermo Fisher Scientific, Waltham, MA, USA). Signals were acquired on the Molecular Imager ChemiDocTM XRS+ Imaging system (BioRad, Neuberg, Germany) and analyzed using the Fiji application (Schindelin et al., 2012). Membranes were stripped using stripping buffer (1 % (w/v) SDS, 0.2 M glycine, pH 2.5), washed 3 times with TBST, blocked and incubated with the second primary antibody, if necessary.

### Proteasome activity assay on MEF cells treated with glyoxal

Proteasome activity assay was performed using the 20S proteasome activity assay kit (Millipore, Billerica, MA, USA) following the manufacturer instructions. Briefly, cell pellets were thawed and ice-cold lysis buffer (50 mM HEPES pH 7.5, 5 mM EDTA, 150 mM NaCl, 1 % (v/v) Triton X-100, 2 mM ATP) was added and left on ice for 30 min with quick vortex steps any 10 min. Samples were centrifuged at 20817 x *g*, for 15 min at 4°C to remove any debris. For protein estimation, a small aliquot was used in order to perform EZQ^®^ Protein Quantitation Kit (Thermo Scientific, Waltham, MA, USA). 50 μg of protein extract were incubated with fluorophore-linked peptide substrate (LLVY-7-amino-4-methylcoumarin [AMC], from the same kit) for 60 min at 37°C. Proteasome activity was measured by quantification of fluorescent units from cleaved AMC at 380/460 nm using a microplate reader m1000 (Tecan). Samples were measured together with positive control supplied with the kit.

### Glyoxalase 1 activity assay

Glyoxalase 1 activity was measured using the ab241019 Glyoxalase I Assay Kit (Colorimetric, Abcam, Cambridge, UK) following manufacturer instructions. Briefly, equal volumes of tissue homogenates (obtained as described above) from heart, liver and kidney, corresponding to an estimated protein amount of 50 μg were analyzed. The substrate mix was prepared and left for 10 min at RT in the darkness to allow the formation of hemimercaptal. Afterwards, samples were transferred to a 96 well-plate and the substrate mix was added. The formation of S-D-lactoylglutathione (SLG) was monitored for 30 min in a kinetic mode by using a microplate reader m1000 (Tecan) at OD 240 nm. The activity was then calculated using the formula suggested by the vendors and normalized for equal loading using immunoblot against beta-actin (as described in “*Immunoblot for mono and polyubiquitinated proteins of MEF cells treated with glyoxal*”) performed on the same homogenates.

### Other data analyses

Gene Ontology enrichment based on ClueGO v2.5.6 (Bindea et al., 2009) was performed using the table S2 (highly confident sites) as input, and by applying the following parameters: for heart (Kappa score threshold 0.4, minimum percentage =5 and number of genes =3), for kidney (Kappa score threshold 0.4, minimum percentage =2 and number of genes =3) and for liver (Kappa score threshold 0.4, minimum percentage =2 and number of genes =3). Statistical test used = Enrichment/depletion (Two-side hypergeometric test). For the overlap between HUVEC cells and MEF cells, the following parameters were applied: Kappa score threshold 0.4, minimum percentage =4 and number of gene =2). Statistical test used = Enrichment/depletion (Two-side hypergeometric test).

KEGG pathway and Gene Ontology enrichments for MEF and HUVEC total proteome were performed by Gene Set Enrichment Analysis (GSEA) with WebGestalt (Liao et al., 2019) using the entire lists of quantified proteins ranked on the basis of the measured log2 fold changes.

### HUVEC lysis and western blot

HUVEC were lysed in ice-cold Tris buffer (50 mM Tris (pH 7.4), 2 mM EDTA, 1 mM EGTA, 50 mM NaF, 10 mM Na_4_P_2_O_7_, 1 mM Na_3_VO_4_, 1 mM DTT, 1 % Triton X-100, 0.1 % SDS, 1 mM PMSF, 10 μl/ml protease inhibitor cocktail) for 15 min on ice, scraped and centrifuged (700 x g, 6 min). Aliquots of supernatants were used for protein determination according to Lowry. Lysates were supplemented with Laemmli buffer, subjected to SDS-PAGE (25-50 μg lysate protein/lane) and blotted onto polyvinylidene difluoride (PVDF) membranes. The membranes were blocked for 1 h in TBST (20 mM Tris (pH 7.6), 137 mM NaCl, 0.1 % (w/v) Tween^®^ 20) containing 5 % non-fat dried skimmed milk. Thereafter, blots were incubated with primary antibodies (diluted in TBST containing 5 % BSA) overnight at 4 °C followed by incubation with horseradish peroxidase-conjugated secondary antibodies for 1 h. Proteins were detected using the enhanced chemiluminescence (ECL) reagent (GE Healthcare, Chicago, IL, USA) or Western Lightning Plus-ECL reagents (Perkin Elmer, Waltham, MA, USA). The intensity of bands was quantified by densitometry using the ImageJ software (NIH, Bethesda, MD, US). Phospho- and acetylation-specific signals were normalized to the signal of the respective total proteins. For evaluation of protein expression, the signals were normalized to β-actin.

### 5-Bromo-2’-deoxyuridine (BrdU) proliferation assay

Glyoxal treatment of HUVEC was performed for 24 h in cell culture dishes and, after reseeding, for another 24 h in 24-well plates. Thereafter, cells were starved in starvation medium (M199, 2 % FCS, 7.5 U/ml heparin, 680 μM glutamine, 100 U/ml penicillin, 100 μg/ml streptomycin) for 4 h and stimulated with bFGF for 24 h. DNA synthesis was analyzed using the Cell Proliferation ELISA, BrdU (colorimetric) kit from Roche Diagnostics (Mannheim, Germany) according to the manufacturer’s protocol. In brief, during the last 5 h of experimental incubation 10 μM BrdU were added. Then, cells were fixed for 30 min in Fix/Denat solution. 200 μl peroxidase-conjugated anti-BrdU-antibody (1:100) were added per well and incubated for 75 min protected from light. Cells were washed three times with PBS before adding 200 μl substrate solution. Reaction was stopped after 5-20 min depending on the color development by adding 50 μl of 1 M H_2_SO_4_ for 1 min. The solution was transferred into a 96-well plate and absorption was measured at 450 nm.

### CFSE proliferation assay

HUVEC were washed twice with warm (37°C) HEPES buffer containing 0.25 % human serum albumin (HEPES/HSA) and incubated with 5 μM CFSE for 15 min at 37 °C. After two washing steps with warm HEPES/FCS, cells were incubated with glyoxal in 1 ml experimental medium for the indicated times. Thereafter, cells were washed twice with PBS, detached with 300 μl trypsin/EDTA and transferred to 700 μl of HEPES/FCS. The cell suspension was pooled with 1 ml HEPES/FCS obtained after rinsing the dish. 10 ml of HEPES buffer were added and samples centrifuged (500 x g, 1 min). Pellets were resuspended in 300 μl PBS and subjected to flow cytometric analysis. Median values were evaluated using the FlowJo™ software (Becton, Dickinson and Company, Ashland, OR, US).

### Angiogenesis assay

Glyoxal treatment was performed for 24 h in 90 mm-cell culture dishes and, during generation of spheroids, for another 24 h in 96-well plates. Spheroids were prepared as previously described (Spengler et al., 2018). In brief, cells suspended in growth medium were mixed at a 4:1 ratio with methyl cellulose (12 mg/ml) and 3000 cells/well were cultured in 96-well round-bottom plates for 24 h. The formed spheroids were collected, centrifuged (200 x g, 4 min) and washed with HEPES buffer. Then, spheroids were transferred to a fibrinogen solution (1.8 mg/ml in HEPES buffer) containing 20 U/ml aprotinin to obtain a suspension with approximately 100 spheroids per ml. 300 μl of this suspension together with 0.2 U thrombin were added per well of a 24-well plate. The plate was incubated for 20 min at 37 °C to allow the formation of a fibrin gel. To equilibrate the gel with medium, M199 containing 2 % FCS, 680 μM glutamine, 100 U/ml penicillin and 100 μg/ml streptomycin was added twice for 15 min. Thereafter, spheroids were cultured in the same medium and stimulated with 50 ng/ml VEGF for 24 h. Finally, spheroids were fixed on ice by adding 1 ml 4 % paraformaldehyde per well for 10 min. After two washing steps with PBS, spheroid sprouting was viewed by light microscopy and pictures were taken (AxioVert 200, Carl Zeiss, Oberkochen, Germany). The number of sprouts was analyzed using cellSens image analysis software (Olympus, Tokyo, Japan).

### Cell synchronization with double thymidine block

Cells were synchronized as described previously (Amin and Varma, 2017). Briefly, one day after seeding 2 mM thymidine was added and incubated with cells for 18 h. After washing (twice with PBS, once with M199), cells were cultured in full growth medium for 9 h and subsequently incubated with 2 mM thymidine for another 18 h. Cells were washed as described and glyoxal was added in experimental medium for the indicated times.

### Cell cycle analysis

During the last 30 min of glyoxal treatment, 10 μM 5-ethynyl-2’-deoxyuridine (EdU) were added to HUVEC cultured on 90 mm-dishes. Cells were washed twice with PBS, detached with 1 ml trypsin/EDTA and transferred to 4 ml of HEPES/FCS. The cell suspension was pooled with 4 ml HEPES/FCS obtained after rinsing the dish and samples were centrifuged. All centrifugations were carried out at 500 x g for 3 min at room temperature if not otherwise stated. Cell pellets were washed once in 1 ml BSA buffer (1 % BSA in PBS), resuspended in 100 μl of the same buffer, fixed with 100 μl of 4 % paraformaldehyde for 15 min, centrifuged and resuspended in 300 μl PBS. 700 μl of 100 % ethanol were added dropwise under constant gentle shaking and samples were frozen overnight. The next day, cells were centrifuged (700 x g, 3 min, RT), washed in 1 ml BSA buffer and permeabilized in 100 μl Triton-based BSA buffer (TBB, 0.2 % Triton X-100 in BSA buffer) for 30 min. Then, cells were centrifuged, resuspended in 500 μl BSA buffer, incubated for 1 h, centrifuged again and incubated in 150 μl of primary antibody solution (1:200 p-H3 (S10) antibody in TBB) for 2 h. This was followed by another addition of 500 μl TBB, centrifugation and incubation of cells in 150 μl of secondary antibody solution (1:500 AF488 goat anti-rabbit antibody in TBB) for 1 h. Next, after adding 500 μl TBB, centrifugation and an additional washing with 1 ml BSA buffer, cells were incubated in 100 μl of Click-iT reaction cocktail (2.5 mM CuSO_4_, 1:200 AF647 azide, 50 mM sodium ascorbate in PBS) for 30 min. Subsequently, another washing with 1 ml BSA buffer was performed and cells were incubated in 300 μl BSA buffer containing 1 μg DAPI for 30 min. Samples were subjected to flow cytometric analysis with triple detection of AF488, AF647 and DAPI. Percentages of cells in the respective cell cycle phases were evaluated using the FlowJo™ software.

### Seahorse analysis of cells

HUVEC were seeded into Seahorse XF96 Cell Culture Microplates (3,000 cells/well; Agilent Technologies, Santa Clara, CA, USA), incubated for 24 h and then treated with glyoxal for 48 h. Mito stress test: Medium was replaced with Seahorse XF Base Medium (103334-100, Agilent Technologies, pH adjusted to 7.4), supplemented with 10 mM D-glucose, 2 mM L-glutamine and 1 mM sodium pyruvate. Cells were then cultured for another hour in a CO_2_-free incubator at 37 °C. Oxygen consumption rates (OCR) and extracellular acidification rates (ECAR) were monitored at basal conditions and after sequential injections of 2 μM oligomycin to block the mitochondrial ATP synthase, 2 μM carbonyl cyanide-4-(trifluoromethoxy)phenylhydrazone (FCCP) to uncouple oxidative phosphorylation and 2 μM antimycin A to fully inhibit mitochondrial respiration. Glycolysis stress test: Medium was replaced with Seahorse XF Base Medium (pH adjusted to 7.4) supplemented with 2 mM L-glutamine, and cells were cultured for another hour in a CO_2_-free incubator at 37 °C. OCR and ECAR were monitored at basal conditions and after sequential injections of 10 mM D-glucose, 2 μM oligomycin and 50 mM 2-deoxy-D-glucose, an inhibitor of glycolysis.

Measurements in both settings were performed in 3 min mix and 3 min measure cycles at 37 °C in six replicates per condition on a Seahorse XFe96 Analyzer (Agilent Technologies). OCR and ECAR were depicted as pmol/min and mpH/min, respectively, and normalized to the exact cell number of each well measured by high-content microscopy. Wave software (Agilent Technologies) was used to analyze the datasets.

### High-content microscopy

Cell supernatants were removed from the Seahorse XF96 microplate and cells were fixed for 10 min with 100 % methanol at room temperature. Cells were then washed once with PBS and incubated for 10 min with 1 μg/ml DAPI at room temperature. After two more washing steps with PBS, cell nuclei were counted on an ImageXpress Micro confocal high-content imaging system (Molecular Devices, San Jose, CA, USA).

### ATP measurements

The intracellular ATP content was determined using the ATP Kit SL from Biotherma (Handen, Sweden) according to the manufacturer’s protocol. At the end of experimental incubations, cell proteins were denaturated by adding 500 μl ethanol per dish. After evaporation of the ethanol, 250 μl Tris buffer of the assay kit was added and one freezing/thawing cycle in liquid nitrogen was performed. Cells were then scraped off, centrifuged (700 x g, 5 min) and supernatants subjected to ATP measurements. For normalization, cells in identically treated dishes were lysed with solubilization buffer (100 mM NaOH, 1.9 M Na_2_CO_3_, 1% SDS) and the protein content was determined according to Lowry.

### Intracellular and mitochondrial ROS measurement

30 min before the end of the indicated glyoxal treatment, HUVEC were washed with PBS, 600 μl of the respective staining solution were added, glyoxal was re-added and incubation was completed. The staining solutions contained 5 μM CM-H2DCFDA or 3 μM MitoSOX™ (both Thermo Scientific, MA, Waltham, USA) diluted in HEPES/HSA for the detection of intracellular or mitochondrial ROS, respectively. Cells were washed with PBS, detached with 300 μl trypsin/EDTA and transferred to 700 μl HEPES/FCS. The cell suspension was pooled with 1 ml HEPES/FCS obtained from rinsing the dish, 10 ml HEPES buffer were added and cells centrifuged (500 x g, 1 min). Cell pellets were resuspended in 300 μl PBS and subjected to flow cytometry analysis. Median values were evaluated using the FlowJo™ software.

### Detection of mitochondrial mass

30 min before the end of the indicated glyoxal treatment HUVEC were washed with PBS and 600 μl of 100 nM MitoTracker™ (Thermo Scientific, Waltham, MA, USA) diluted in HEPES/HSA were added. Glyoxal was re-added and incubation was completed. Cells were washed with PBS, detached with 300 μl trypsin/EDTA and transferred to 700 μl HEPES/FCS. The cell suspension was pooled with 1 ml HEPES/FCS obtained from rinsing the dish, 10 ml HEPES/HSA were added and cells centrifuged (500 x g, 1 min). Pellets were resuspended in 300 μl PBS and subjected to flow cytometry analysis. FlowJo™ software was used to evaluate the median values of MitoTracker™-positive cells

### SA-β-Gal staining

After glyoxal incubation, cells were washed twice with 1 ml cold PBS and fixed with 1 ml of a 2 % formaldehyde/0.2 % glutaraldehyde solution for 3 min. SA-β-Gal staining was performed as previously described (Debacq-Chainiaux et al., 2009). Briefly, after two washing steps with PBS, 1.5 ml staining solution (40 mM citric acid/Na phosphate buffer, 5 mM K_4_[Fe(CN)_6_]•3H_2_O, 5 mM K_3_[Fe(CN)_6_], 150 mM NaCl, 2 mM MgCl_2_, 1:20 X-Gal (20 mg/ml in DMF) in water, pH 6.0) was incubated with cells at 37 °C overnight. Pictures were taken (EVOS™ FL Auto, Thermo Scientific, Waltham, MA, USA) and the number of SA-β-Gal-positive cells was counted using ImageJ and normalized to the total cell number.

### γH2A.X (S139) immunofluorescence staining

HUVEC were washed with warm (37 °C) HEPES buffer, fixed in ice-cold 4 % paraformaldehyde for 15 min and washed twice with PBS. Cells were permeabilized in PBS containing 0.3 % Triton X-100 for 5 min. Blocking solution (1 % BSA-C, 5 % goat serum in PBS) was added for 1 h. After two washing steps with PBS, samples were incubated in γH2A.X (S139) antibody diluted in blocking solution (1:400) in a humidified chamber overnight at 4 °C. Cells were washed twice with PBS and incubated with AF488-labelled goat anti-rabbit secondary antibody (1:500 in blocking solution) for 2 h. After two washing steps with PBS, DAPI (1 μg/ml in PBS) was added for 10 min and cells were washed three times with PBS before mounting the coverslips on microscopic slides using Fluoromount-G.

### α/β-Tubulin and ac-Tubulin (K40) immunofluorescence staining

Washing, fixation and permeabilization of HUVEC were performed as described for γH2A.X (S139) staining. Thereafter, cells were incubated for 2 h with primary antibodies against α/β-tubulin or ac-tubulin (K40) diluted in blocking solution (1:100). After two washing steps with PBS, an AF488-labelled goat anti-rabbit secondary antibody (1:500 in blocking solution) was added for 1 h. Cells were washed three times with PBS. Then, DAPI (1 μg/ml in PBS) was added for 10 min and cells were washed three times with PBS before mounting the coverslips on microscopic slides using Fluoromount-G. For immunofluorescence, a LEICA DMi8 TCS SP8 inverted laser scanning microscope was employed (Leica Biosystems, Wetzlar, Germany) and pictures were taken. Evaluation was performed via a blinded approach, in which cells with strongly structured tubulin filaments per high-power field were counted and normalized to the number of DAPI stained nuclei.

### Data availability

The mass spectrometry proteomics data have been deposited to the ProteomeXchange Consortium via the PRIDE (Perez-Riverol et al., 2019) partner repository with the dataset identifiers:

TMT-10plex data for organ aging proteome: PXD021883
DIA for acetylated peptides: PXD021891
MQ searches for CMLpepIP: PXD021921
DIA for MEF and HUVEC cells: PXD021985

In addition, the proteomics data presented in this manuscript are available via a R shiny web server: https://genome.leibniz-fli.de/shiny/orilab/CMLsites/

