## Supplemental information for "Mapping sites of carboxymethyllysine modification on proteins reveals its consequences for proteostasis and cell proliferation"

**This PDF file includes:**

Figures S1 to S6

**The following supplemental tables are provided as separate Excel files:**

Table S1: List of CML sites identified in MEF and HUVEC. Related to Figure 1 and S1.

Table S2: List of CML sites identified in mouse organs and proteome changes induced by aging. Related to Figure 2 and S2.

Table S3: Proteome changes induced in MEF by glyoxal treatment. Related to Figure 3 and S3.

Table S4: Proteome changes induced in HUVEC by glyoxal treatment. Related to Figure 4.

Table S5: List of cell cycle-related proteins affected in HUVEC by glyoxal or directly modified by CML. Related to Figure 5.

### Supplemental Figures

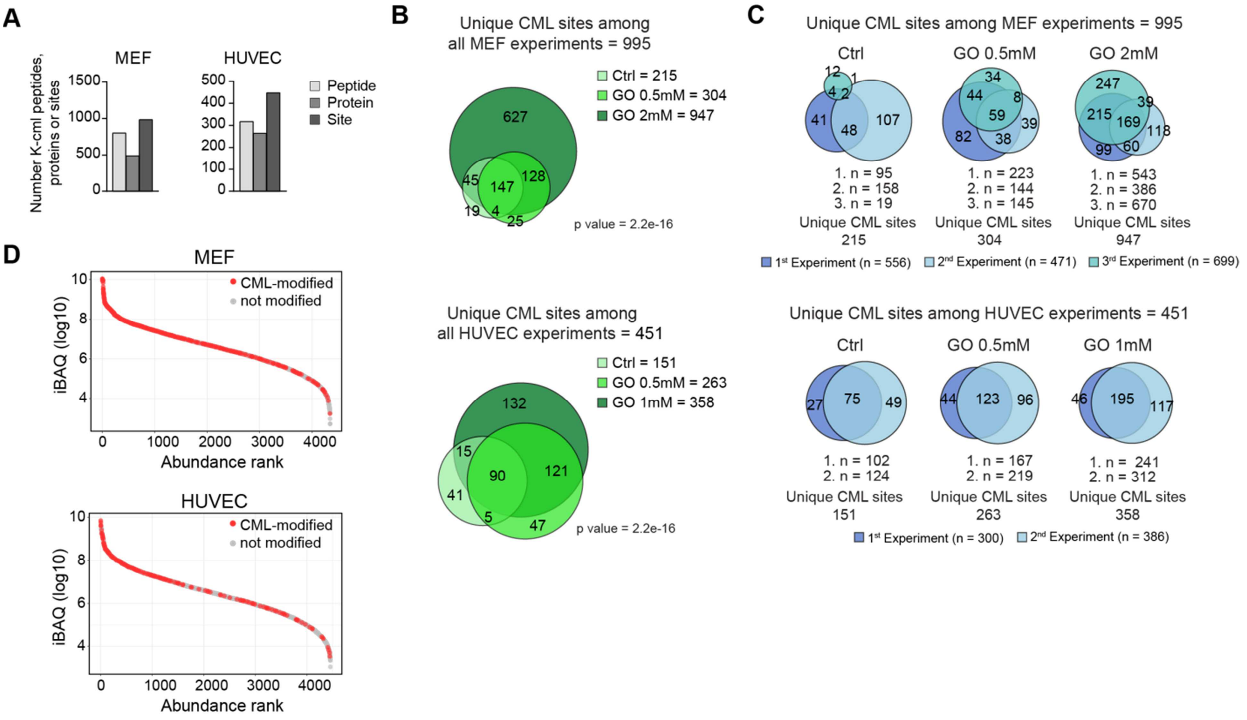


**Figure S1.**
**A.** Statistics of CML sites identified in MEF and HUVEC treated with glyoxal. Data from different treatments and biological replicates were combined. **B.** Overlap between CML sites identified under different experimental conditions. **C.** Reproducibility of CML site identification across independent experiments. In B and C, only highly confident sites were considered and the overlap between experiment was tested using Fisher's exact test. **D.** Rank plot showing the abundance distribution of proteins identified as targets of CML. Related to Figure 1 and Table S1.

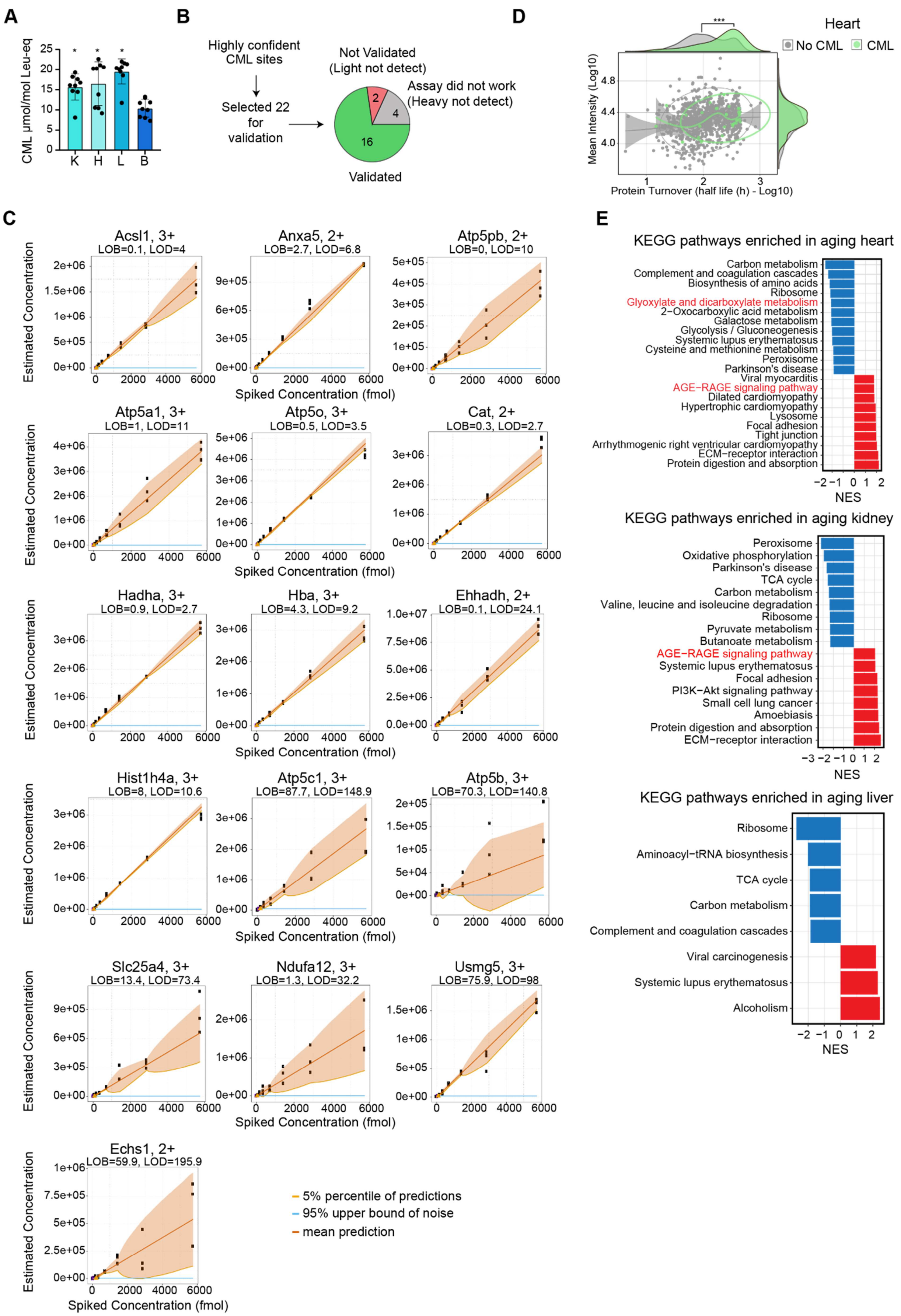
A
B
D
CML µmol/mol Leu-eq
K
H
L
B
Highly confident CML sites
Selected 22 for validation
Not Validated (Light not detect)
Assay did not work (Heavy not detect)
Validated
Heart
No CML
CML
Mean Intensity (Log10)
Protein Turnover (half life (h) - Log10)
C
Acsl1, 3+
LOB=0.1, LOD=4
Anxa5, 2+
LOB=2.7, LOD=6.8
Atp5pb, 2+
LOB=0, LOD=10
Atp5a1, 3+
LOB=1, LOD=11
Atp5o, 3+
LOB=0.5, LOD=3.5
Cat, 2+
LOB=0.3, LOD=2.7
Hadha, 3+
LOB=0.9, LOD=2.7
Hba, 3+
LOB=4.3, LOD=9.2
Ehhadh, 2+
LOB=0.1, LOD=24.1
Hist1h4a, 3+
LOB=8, LOD=10.6
Atp5c1, 3+
LOB=87.7, LOD=148.9
Atp5b, 3+
LOB=70.3, LOD=140.8
Slc25a4, 3+
LOB=13.4, LOD=73.4
Ndufa12, 3+
LOB=1.3, LOD=32.2
Usmg5, 3+
LOB=75.9, LOD=98
Echs1, 2+
LOB=59.9, LOD=195.9
Estimated Concentration
Spiked Concentration (fmol)
5% percentile of predictions
95% upper bound of noise
mean prediction
E
KEGG pathways enriched in aging heart
Carbon metabolism
Complement and coagulation cascades
Biosynthesis of amino acids
Ribosome
Glyoxylate and dicarboxylate metabolism
2−Oxocarboxylic acid metabolism
Galactose metabolism
Glycolysis / Gluconeogenesis
Systemic lupus erythematosus
Cysteine and methionine metabolism
Peroxisome
Parkinson's disease
Viral myocarditis
AGE−RAGE signaling pathway
Dilated cardiomyopathy
Hypertrophic cardiomyopathy
Lysosome
Focal adhesion
Tight junction
Arrhythmogenic right ventricular cardiomyopathy
ECM−receptor interaction
Protein digestion and absorption
NES
KEGG pathways enriched in aging kidney
Peroxisome
Oxidative phosphorylation
Parkinson's disease
TCA cycle
Carbon metabolism
Valine, leucine and isoleucine degradation
Ribosome
Pyruvate metabolism
Butanoate metabolism
AGE−RAGE signaling pathway
Systemic lupus erythematosus
Focal adhesion
PI3K−Akt signaling pathway
Small cell lung cancer
Amoebiasis
Protein digestion and absorption
ECM−receptor interaction
NES
KEGG pathways enriched in aging liver
Ribosome
Aminoacyl−tRNA biosynthesis
TCA cycle
Carbon metabolism
Complement and coagulation cascades
Viral carcinogenesis
Systemic lupus erythematosus
Alcoholism
NES

1

2 **Figure S2.**

**A.** Quantification of total CML levels in mouse organs. K: kidney, H: heart, L: liver, B: brain. n=9. * $p<0.05$ vs. brain, ordinary one-way ANOVA. **B.** Validation of identified CML sites using parallel reaction monitoring (PRM) and heavy labelled spiked-in reference peptides. **C.** Limit of detection (LOD) and limit of blank (LOB) for PRM assays targeting selected CML sites. Dilution series of synthetic heavy peptides were used for LOB and LOD estimation (see Methods for details). **D.** Protein turnover of CML-modified proteins grouped according to their subcellular localization, as defined by Gene Ontology annotation. Turnover data were taken from (Fornasiero et al., 2018). C: cytoplasm; ER: endoplasmic reticulum; GA: Golgi apparatus; M: mitochondria; N: nucleus. Scatterplot comparing protein abundance (average protein intensity) and protein turnover (half life) of CML-modified proteins. Density plot shows the distribution of each dataset. n=5. *** $p<0.001$, Wilcoxon Rank Sum test with continuity correction. **E.** KEGG pathway Gene Set Enrichment Analysis (GSEA) in aging tissues performed using WebGestalt. Normalized enrichment score (NES) indicates pathways enriched among proteins that increase (red) or decrease (blue) with aging (FDR<0.05). All the proteins quantified in each experiment were ranked according to their log2 fold change and used as input for GSEA. Related to Figure 2 and Table S2.

**A**

| ID | Gene | Type | Protein name | Where (CML site) |
|---|---|---|---|---|
| Q9R1P1 | PSMB3 | 26S Proteasome | Proteasome subunit beta type-3 | Liver (15) |
| Q9R1P4 | PSMA1 | 26S Proteasome | Proteasome subunit alpha type-1 | MEF (30) |
| Q8C7R4 | UBA6 | E1 ligase | Ubiquitin-like modifier-activating enzyme 6 | MEF (519) |
| Q9Z1F9 | UBA2 | E1 ligase | SUMO-activating enzyme subunit 2 | MEF (271), HUVEC (271) |
| Q6ZPJ3 | UBE2O | E2 ligase | E2/E3 hybrid ubiquitin-protein ligase UBE2O | Kidney (885, 888), MEF (368, 379, 380, 382, 360) |
| Q61187 | TSG101 | E2 ligase | Tumor susceptibility gene 101 protein | Kidney (9, 10) |
| P68037 | UBE2L3 | E2 ligase | Ubiquitin-conjugating enzyme E2 L3 | MEF (73) |
| Q69ZR2 | HECTD1 | E3 ligase | E3 ubiquitin-protein ligase HECTD1 | Kidney (2285, 2290) |
| Q7TMY8 | HUWE1 | E3 ligase | E3 ubiquitin-protein ligase HUWE1 | MEF (4137, 4142, 3257, 3274), HUVEC (4134, 4139) |
| P46935 | NEDD4 | E3 ligase | E3 ubiquitin-protein ligase NEDD4 | MEF (186, 429, 484) |
| Q8BFU3 | RNF214 | E3 ligase | RING finger protein 214 | MEF (136, 142) |
| P58283 | RNF216 | E3 ligase | E3 ubiquitin-protein ligase RNF216 | MEF (629) |
| Q8K2Y0 | RNF219 | E3 ligase | RING finger protein 219 | MEF (498, 505) |
| Q62318 | TRIM28 | E3 ligase | Transcription intermediary factor 1-beta | MEF (770, 774, 779), HUVEC (366, 304) |
| Q6WKZ8 | UBR2 | E3 ligase | E3 ubiquitin-protein ligase UBR2 | MEF (1496) |
| A2AN08 | UBR4 | E3 ligase | E3 ubiquitin-protein ligase UBR4 | MEF (1929), HUVEC (1060, 1931, 1141, 1150) |
| Q0P5W1 | VPS8 | E3 ligase | Vacuolar protein sorting-associated protein 8 homolog | MEF (1313, 1316) |
| Q80YR4 | ZNF598 | E3 ligase | Zinc finger protein 598 | MEF (94) |
| Q9Y4D8 | HECTD4 | E3 ligase | Probable E3 ubiquitin-protein ligase HECTD4 | HUVEC (2326, 2331) |
| P29590 | PML | E3 ligase | Protein PML | HUVEC (487, 490) |
| P0CG49 | UBB | Ubiquitin | Polyubiquitin-B | Heart (11), HUVEC (11, 63), MEF (6, 11, 63) |
| P0CG50 | UBC | Ubiquitin | Polyubiquitin-C | Heart (11), HUVEC (11, 63), MEF (6, 11, 63) |
| P62983 | RPS27A | Ubiquitin | Ubiquitin-40S ribosomal protein S27a | Heart (11), HUVEC (11, 63), MEF (6, 11, 63) |
| P62984 | UBA52 | Ubiquitin | Ubiquitin-60S ribosomal protein L40 | Heart (11), HUVEC (11, 63), MEF (6, 11, 63) |
| Q99LD4 | GPS1 | UPS associated | COP9 signalosome complex subunit 1 | MEF (204) |
| P56399 | USP5 | UPS associated | Ubiquitin carboxyl-terminal hydrolase 5 | MEF (288, 291), HUVEC (288, 291) |
| P35125 | USP6 | UPS associated | Ubiquitin carboxyl-terminal hydrolase 6 | HUVEC (313) |

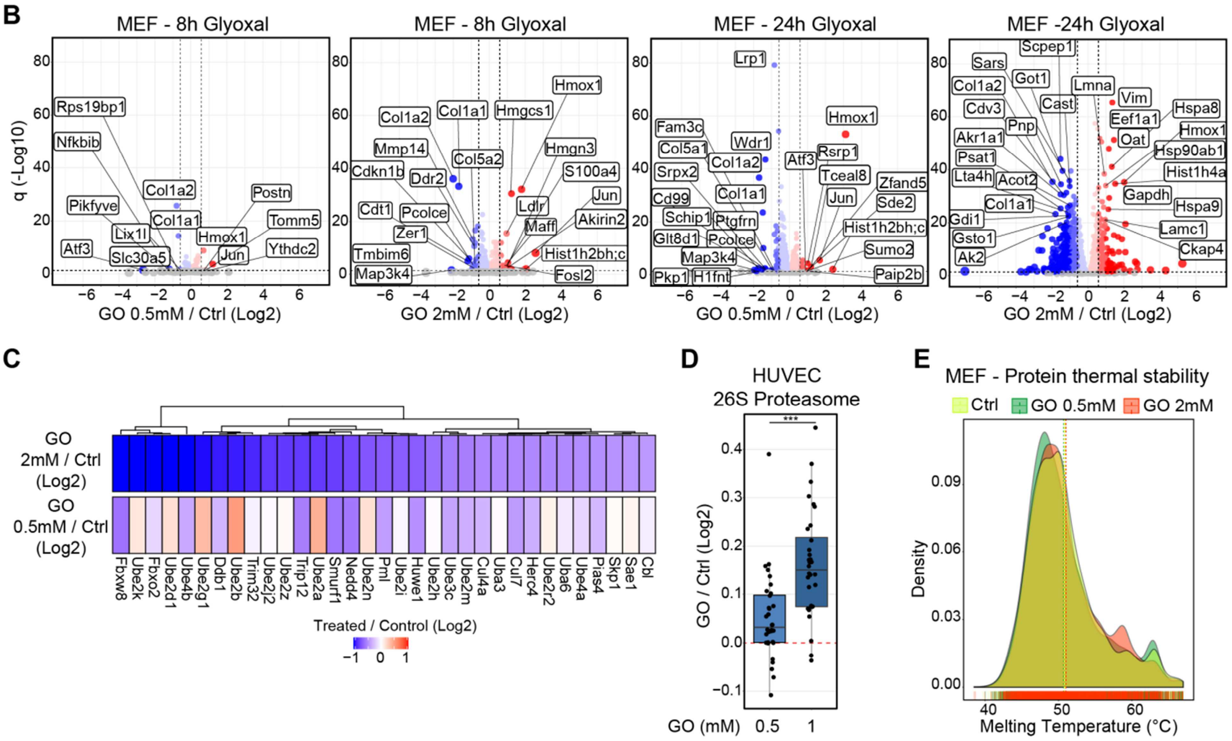


**Figure S3.**
**A.** Table of CML sites identified on proteins related to the ubiquitin proteasome system. **B.** Protein abundance changes induced in MEF treated with glyoxal (GO). Volcano plots depict proteins that significantly increase (red) or decrease (blue) abundance upon GO treatment. Proteins not affected are shown in gray. Horizontal dashed line indicates a significance cut-off of $q<0.05$ and vertical dashed lines an absolute fold change (log2)>0.58. **C.** Heatmap representing the protein abundance of protein involved in the ubiquitin-mediated proteolysis (Fig. 3D in red) induced in MEF by 0.5 mM and 2 mM GO after 24 h of treatment. **D.** Boxplot of fold changes for members of the 26S proteasome in HUVEC cells treated with GO for 48 h compared to control (Ctrl). n=4. *** $p<0.001$, Wilcoxon Rank Sum test with continuity correction. **E.** Effect of GO on protein thermal stability in MEF treated for 24 h. Melting temperatures for 4367 (Ctrl), 4174 (GO 0.5 mM) and 4646 (GO 2 mM) protein groups were estimated using thermal proteome profiling (see Methods for details). Related to Figure 3 and Table S3.

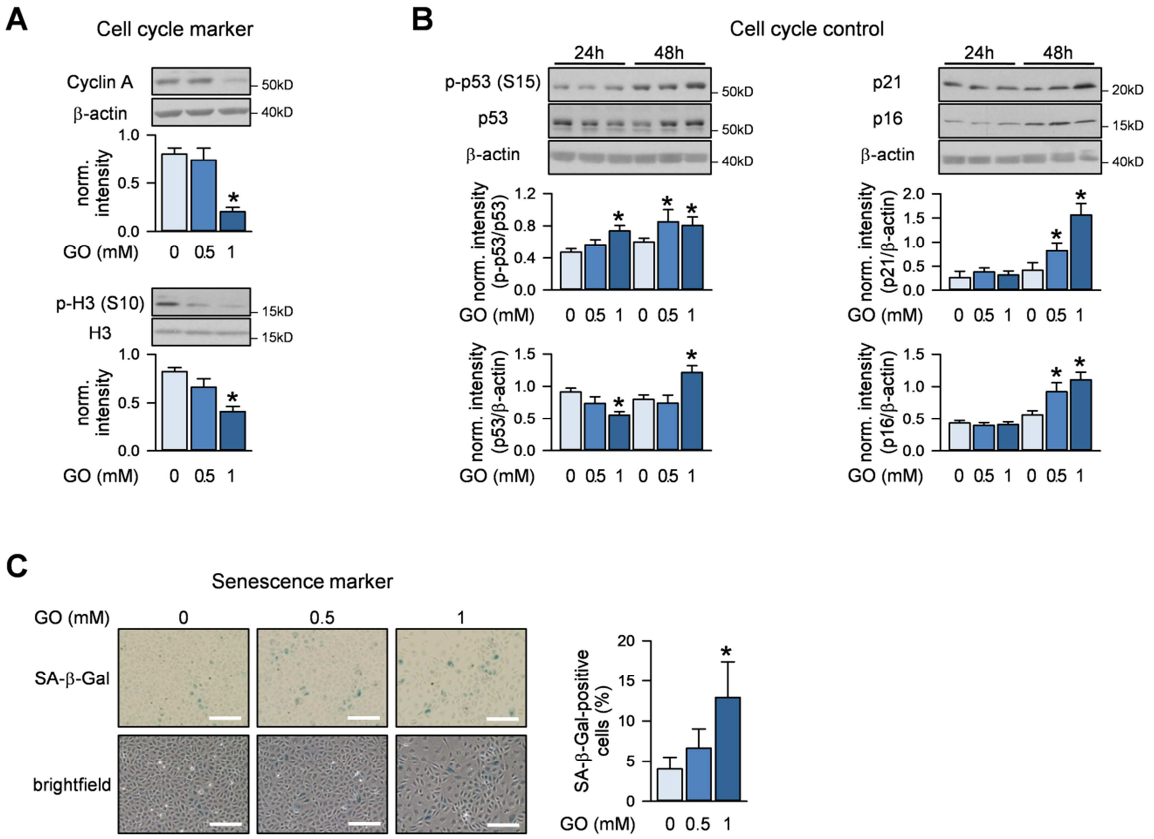


**Figure S5.**
**A-C.**: HUVEC were treated with GO at the indicated concentrations for 24 h (**A,B**) or 48 h (**B,C**). **A-B**. Cell lysates were subjected to immunoblot analysis of cell cycle regulators. n=6 (**A**), n=5 (**B**), * p<0.05 vs. respective control. **C**. Senescence-associated β-galactosidase (SA-β-Gal) was stained. Representative pictures and quantification of SA-β-Gal-positive cells are shown. n=5, * p<0.05 vs. control. Statistical significance was analyzed using one-way repeated measurement ANOVA corrected using Holm-Šidák method. Related to Figure 5.

| ID | Gene | Protein name | Where (CML site) |
|---|---|---|---|
| P05213 | Tubulin alpha-1B chain | TUBA1B | MEF (336, 338, 326, 394, 401, 163, 164, 96, 352, 370, 60), HUVEC (394, 401, 336, 338, 163, 164) |
| P68369 | Tubulin alpha-1A chain | TUBA1A | MEF (336, 338, 326, 394, 401, 163, 164, 96, 60, 352, 370), HUVEC (394, 401, 326, 336, 338, 163, 164) |
| P05214 | Tubulin alpha-3 chain | TUBA3A | MEF (336, 338, 326, 394, 401, 163, 164, 96, 60, 352, 370) |
| P68373 | Tubulin alpha-1C chain | TUBA1C | MEF (336, 338, 326, 394, 401, 163, 164, 96, 60, 352, 370), HUVEC (394, 401, 326, 336, 338, 163, 164) |
| P68368 | Tubulin alpha-4A chain | TUBA4A | MEF (394, 401, 163, 352, 370), HUVEC (394, 401, 163, 164) |
| Q9JJZ2 | Tubulin alpha-8 chain | TUBA8 | MEF (394, 401, 96, 352, 370) |
| P68372 | Tubulin beta-4B chain | TUBB4B | MEF (58, 379, 324, 336, 103, 216), Kidney (324) |
| P99024 | Tubulin beta-5 chain | TUBB5 | MEF (362, 379, 58, 324, 336, 103, 216), Kidney (324) |
| Q922F4 | Tubulin beta-6 chain | TUBB6 | MEF (57, 58) |
| Q3UX10 | Tubulin alpha chain-like 3 | TUBAL3 | MEF (401), HUVEC (401, 408) |
| Q7TMM9 | Tubulin beta-2A chain | TUBB2A | MEF (379, 324, 336, 103, 216), HUVEC (324, 103, 379), Kidney (324) |
| Q9CWF2 | Tubulin beta-2B chain | TUBB2B | MEF (379, 324, 336, 103, 216), HUVEC (324, 103, 379), Kidney (324) |
| Q9D6F9 | Tubulin beta-4A chain | TUBB4A | MEF (103, 216), HUVEC (58, 324, 103) |
| Q13748 | Tubulin alpha-3C/D chain | TUBA3C | HUVEC (394, 401, 326, 336, 338, 163, 164) |
| P07437 | Tubulin beta chain | TUBB | HUVEC (58, 379, 324, 103) |
| Q6PEY2 | Tubulin alpha-3E chain | TUBA3E | HUVEC (326, 336, 338) |
| Q13509 | Tubulin beta-3 chain | TUBB3 | HUVEC (58) |
| P04350 | Tubulin beta-4A chain | TUBB4A | HUVEC (103) |

**Figure S6.**
Table of CML sites identified on tubulins. Related to Figure 6.
